# Intercellular communication atlas reveals Oprm1 as a neuroprotective factor for retinal ganglion cells

**DOI:** 10.1101/2023.07.14.549118

**Authors:** Cheng Qian, Ying Xin, Cheng Qi, Hui Wang, Bryan C. Dong, Donald Zack, Seth Blackshaw, Samer Hattar, Feng-Quan Zhou, Jiang Qian

## Abstract

The progressive death of mature neurons often results in neurodegenerative diseases. While the previous studies have mostly focused on identifying intrinsic mechanisms controlling neuronal survival, the extracellular environment also plays a critical role in regulating cell viability. Here we explore how intercellular communication contributes to the survival of retinal ganglion cells (RGCs) following the optic nerve crush (ONC). Although the direct effect of the ONC is restricted to the RGCs, we observed transcriptomic responses in other retinal cells to the injury based on the single-cell RNA-seq, with astrocytes and Müller glia having the most interactions with RGCs. By comparing the RGC subclasses with distinct resilience to ONC-induced cell death, we found that the high-survival RGCs tend to have more ligand-receptor interactions with other retinal cells, suggesting that these RGCs are intrinsically programmed to foster more communication with their surroundings. Furthermore, we identified the top 47 interactions that are stronger in the high-survival RGCs, likely representing neuroprotective interactions. We performed functional assays on one of the receptors, μ-opioid receptor (Oprm1), a receptor known to play roles in regulating pain, reward, and addictive behavior. Although Oprm1 is preferentially expressed in intrinsically photosensitive retinal ganglion cells (ipRGC), its neuroprotective effect could be transferred to multiple RGC subclasses by selectively overexpressing Oprm1 in pan-RGCs in ONC, excitotoxicity, and glaucoma models. Lastly, manipulating Oprm1 activity improved visual functions or altered pupillary light response in mice. Our study provides an atlas of cell-cell interactions in intact and post-ONC retina, and a strategy to predict molecular mechanisms controlling neuroprotection, underlying the principal role played by extracellular environment in supporting neuron survival.

## INTRODUCTION

Neuronal cell death can lead to irreversible loss of sensory, motor, and cognitive functions (*1, 2*). Enhancing neuroprotection is therefore one of the major strategies for potentially delaying the development of neurological diseases. While previous studies have identified many genes and signaling pathways governing neuronal cell death and/or neuroprotection (*3-6*), most attention has been focused on the role of intrinsic factors and cell-autonomous regulatory mechanisms. However, tissues contain diverse cell types that collectively form and regulate the microenvironment (*7, 8*). Although the importance of the microenvironment has been well recognized and extensively investigated in context of stem cell biology, immunology, and cancer fields (*9-11*), it is much less clear how it functions in the mature nervous system to regulate diverse neurological functions.

One major form of intercellular communications in the tissue microenvironment is ligand-receptor interactions, which potentially includes autocrine, paracrine, and endocrine signaling. In the retina, there are many types of cells, such as retinal ganglion cells (RGCs), amacrine cells, bipolar cells, Müller glia, astrocytes, microglia, photoreceptors, and epithelial cells (*12*). Cell-cell interactions in the retina are not only critical for retinal circuits formation during development and maintaining normal retinal functions in adult animals, but also regulate tissue repair after injury. For instance, during retinal development, amacrine cells can regulate RGC maturation and their intrinsic axon growth ability via direct cell-cell contact (*13*). In adult mammals, microglia have been shown to limit the spontaneous neurogenic ability of Müller glia via secretion of TNF-α (*14*). However, these studies mainly focus on interactions between two cell types via limit number of interactions. Recent advances in single-cell sequencing technologies provide the opportunity to systematically map cell-cell interactions based on the expression of ligands, receptors, and other associated genes in each cell type (*15, 16*).

In this study, we employed the optic nerve crush (ONC) model, which exhibits similarities in gene expression patterns with glaucoma (*17*) and is a widely used model to investigate neuronal cell death and survival in the central nervous system *(3-5)*. This model is particularly suitable for studying cell-cell communication since the physical injury specifically affects RGCs, and the alterations in other retinal cells are therefore likely to be triggered through cell-cell communications with RGCs. To comprehensively explore the cellular effects of ONC, we conducted single-cell sequencing of all retinal cells at various time points after the procedure. By doing so, we constructed an atlas of cell-cell communication that systematically documents the identities of individual ligand-receptor interactions and their dynamic regulation in response to ONC. Moreover, through a comparative analysis of interactions among different RGC subclasses, each exhibiting varying levels of resilience to cell death, we successfully identified numerous potential neuroprotective ligand-receptor interactions between the RGCs and neighboring cells.

We identified the μ-opioid receptor, encoded by the *Oprm1* gene, as a novel neuroprotective factor for RGCs. The μ-opioid receptor is the first discovered opioid receptor, and plays a central role in regulating pain, reward, and addictive behaviors (*18-20*). We functionally validated Oprm1 as a neuroprotective factor for RGCs, by demonstrating that overexpression of Oprm1 in RGCs not only led to a markedly increased survival rate of RGCs following several different types of retinal injury, but also significantly improved visually-guided perception behavior. Our study establishes an effective strategy to identify functionally important cell-cell interactions in a complex tissue microenvironment.

## RESULTS

### Effect of optic nerve crush on retinal cell transcriptomes

We performed droplet-based single-cell RNA sequencing (scRNA-seq) on cells dissociated from the mouse whole retina collected either under sham condition or at three time points after the optic nerve crush (ONC) (Fig. 1A). In total, we profiled 56,531 retinal cells, which were clustered and annotated as 14 major cell types based on the known retinal cell type markers (Fig. 1, B to D). For example, *Abca8a*, *Rpe65*, *and Opn1mw* are representative marker genes for Müller glia, retinal pigment epithelial cells, and cone cells, respectively (Fig. 1C). The specific expression patterns of the known marker genes demonstrated the high quality of the cell annotation (Fig. 1D).

**Fig. 1.**
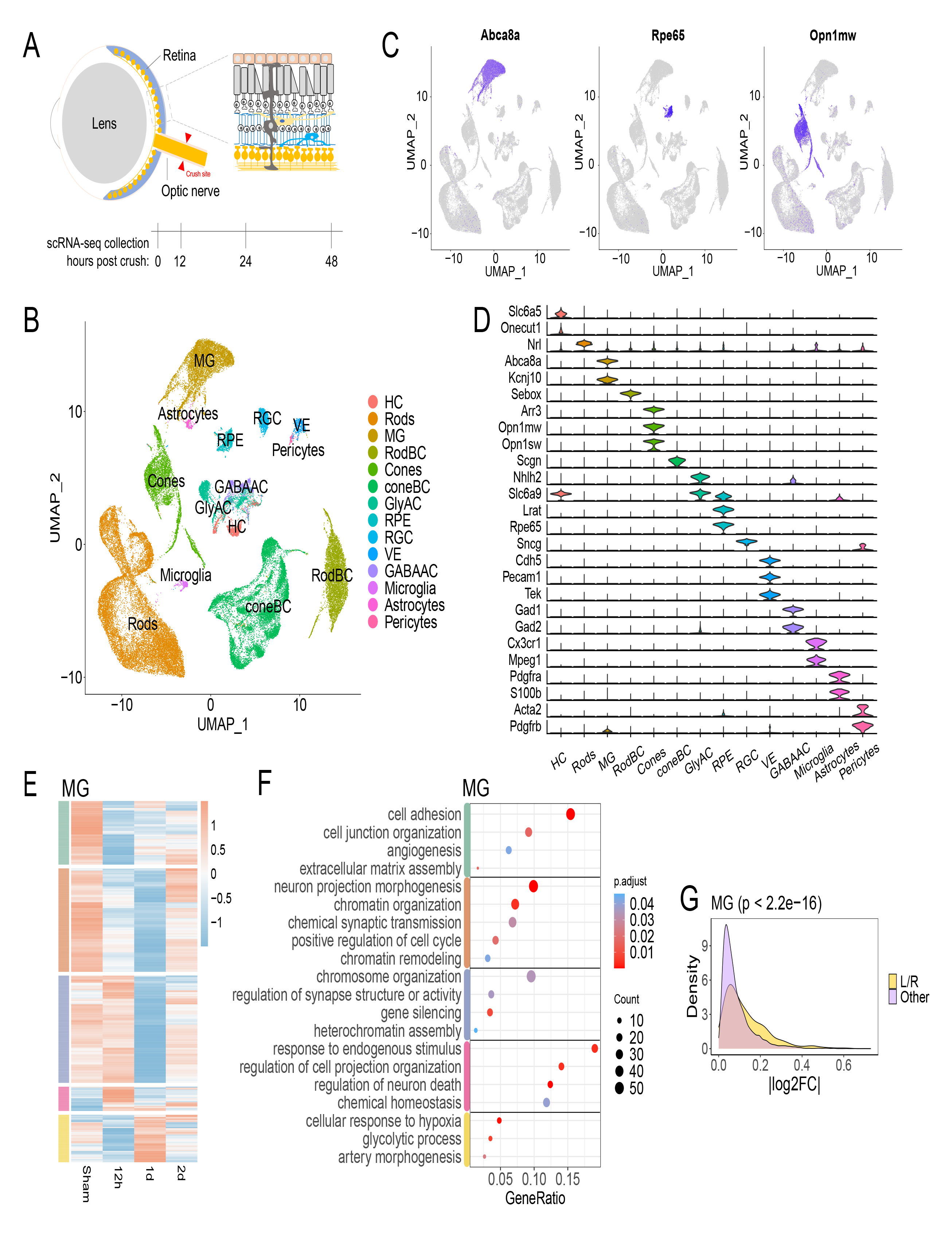
Retinal cells in response to optic nerve crush. (**A**) Schematic diagram illustrates the crush site on the optic nerve, and the timeline for retinal cells being dissociated and collected following the optic nerve crush (ONC). (**B**) UMAP exhibits cell types identified from scRNA-seq data of whole mouse retinal cells before and after ONC. A total of 56,531 retinal cells were used for the analysis and 14 major cell types were identified. (**C**) UMAP shows the expression pattern of *Abca8a, Rpe65* or *Opn1mw* in whole mouse retinal cells, which represent the marker genes for Müller glia (MG), retinal pigment epithelium (RPE) cells and cone cells, respectively. (**D**) Expression patterns of representative known marker genes in major retinal cell types. (**E**) Heatmap exhibits differentially expressed genes (DEGs) patterns in the Müller glia (MG), as an example, at different time points before and after ONC. (**F**) Gene ontology analysis (GO) reveals representative biological processes based on DEG pattern shown in panel (E). (**G**) Density plot illustrates the absolute fold-change (log2FC) of genes expressed in MG as one example (detection rate > 0.1). Ligand and receptor genes (L/R) shown with yellow color are grouped together, while the other genes are shown with light purple for comparison.

We first asked if the ONC affected the gene expression in non-RGC retinal cells. To address this question, we compared the gene expression patterns in each retinal cell type before and after the ONC. The results showed that all identified retinal cell types had differential expressed genes (DEGs) in response to ONC with distinct patterns at different time points (Fig. 1E and fig. S1). For instance, in the Müller glia cells (MG), ONC led to down and up-regulation of multiple sets of genes with different time courses (Fig. 1E). We then performed the Gene Ontology (GO) analysis of the genes with altered expression levels in retinal cell types (Fig. 1F and fig. S1). Interestingly, genes associated with chromatin remodeling were transiently downregulated in MG (Fig. 1F), in line with their inability to be reprogrammed after the retinal injury (*21*). In addition, the genes associated with cytokine production and immune response were enriched in microglia (fig. S1B), consistent with their roles as retinal immune cells. Moreover, genes involved in protein translation and glycolytic metabolism were upregulated in almost all retinal cell types (Fig. 1E and fig. S1), suggesting that retinal cells shift energy production modes and cellular state in response to the surroundings (*22*). The reason for such changed gene transcription in non-RGC cell types is likely through ligands expressed from RGCs in response to ONC. Indeed, genes encoding ligands and receptors as a group are more likely to be differentially expressed than genes of all functional categories (Fig. 1G and fig. S1).

### Ligand-receptor interactions induced by ONC

To identify individual ligand-receptor interactions between RGCs and other retinal cell types, we integrated our scRNA-seq dataset with a publicly available dataset for purified RGCs, obtained at the same time points post-ONC (12h, 24h, and 48h) (*4*). The mean gene expression profile of 1,055 RGC cells from our collection, and that of 86,426 RGCs obtained by Tran et al., (*4*) exhibited a high degree of correlation (fig. S2A). This indicates that both datasets are compatible and the ONC procedures established in different labs were comparable. For subsequent analysis of cell-cell communications between RGCs and other retinal cells, we used the scRNA-seq dataset from Tran et al. for RGCs, which encompasses the majority of known RGC subclasses.

We next used LRLoop, our recently developed computational method for analyzing ligand-receptor based cell-cell communications, to explore intercellular interactions between RGCs and every other retinal cell type (*23*). This method utilized the transcriptome data of both the sender and receiver cells, adjusting interaction scores based on the presence of ligand-receptor feedback loops between the cell types. LRLoop was first used to identify initial signaling interactions from RGCs to other retinal cells, which are termed as “distress” interactions.

Several key characteristics were associated with these distress interactions. First, while all cell types were receivers of distress signals, based on the number of ligand-recpetor pairs, the strongest interactions were seen in astrocytes, Müller glia, GABAergic amacrine, and vasculature pericytes (Fig. 2A). The number of interactions is largely correlated with their physical proximity to the Ganglion Cell Layer (GCL). For instance, astrocytes and Müller glia, in close contact with the GCL (*12*), received the most ligand-receptor interactions from RGCs. Second, most ligand-receptor interactions were common across cell types. Specifically, any ligand produced by RGCs can be detected by most retinal cells with the corresponding receptor (Fig. 2A). However, a few distress signals were specific to one cell type (Fig. 2B), in which microglia were projected to receive the most unique interactions, in line with their role as resident inflammatory cells (*24*) (Fig. 2, A and B). Third, the distress ligand-receptor interactions displayed both transient and prolonged dynamics post-ONC (Fig. 2C). As an example, interactions from RGCs to astrocytes could be grouped into four primary categories based on the similarity of the dynamic interaction patterns (Fig. 2C). The first two groups responded transiently to the injury, being either repressed or activated, but mostly returning to their original states one day post-ONC. The last two groups, in contrast, displayed a quick but sustained or a slowly activated interactions. Finally, the functional roles associated with these 4 types of interaction groups showed overlap, including axonogenesis, neuron projection guidance, and axon guidance. (Fig. 2D).

**Fig. 2.**
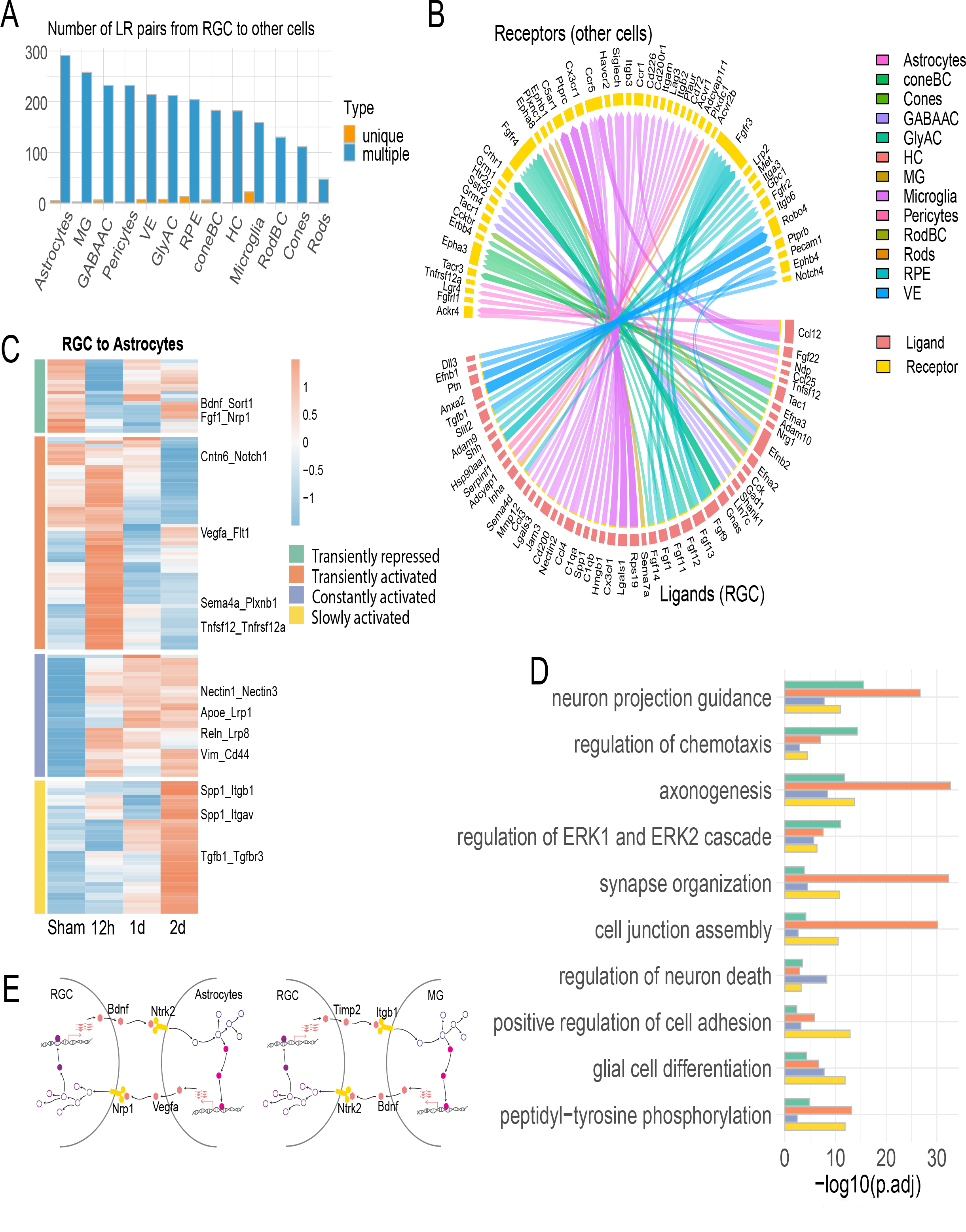
Distress interactions from RGCs to other retinal cells. (**A**) The number of ligand-receptor interactions identified from RGCs to other retinal cells. Interactions identified in multiple and unique cell types are colored dark blue and orange, respectively. (**B**) The specific ligand-receptor interactions identified from RGCs to a unique retinal cell type, which are shown in orange color columns in the panel (A). Genes in the bottom half of the circle are the ligands secreted from RGCs, and the genes in the top half of the circle are the receptors in other retinal cells. The colors of the connecting edges represent the receiver cell types. (**C**) Heatmap reveals the ONC-induced temporal patterns of variable interactions from RGCs to astrocytes, as one example, across time points (the fold change (FC) of the interaction scores between any two time points > 1.2). Four major groups were identified based on their dynamic patterns. (**D**) Gene Ontology (GO) analysis reveals representative biological processes of variable ligand-receptor interactions in each group identified in the panel (C). The enrichment was calculated with all expressed genes (detection rate > 0.1) in either RGCs or astrocytes in any time point as the background. (**E**) Examples of cell-cell feedback loops. A ligand from the sender cell interacts with a receptor on the receiver cell, triggering gene transcription in receiver cells, in which some ligand genes are expressed and sent back to the original sender cells.

We subsequently identified the reciprocal “responsive” ligand-receptor interactions, sent from the other retinal cells back to RGCs. Based on our bioinformatics analyses, some of these responsive interactions might be triggered by the distress interactions, thereby forming a cell-cell signaling loop (Fig. 2E and fig. S2B). For instance, VEGF-A is known to be neuroprotective for RGCs under stress (*25, 26*). Our analysis showed that secretion of the neuroprotective VEGF-A by astrocytes could potentially be initiated by the Bdnf-Ntrk2 distress interaction from RGCs to astrocytes, as these interactions formed a loop through signaling and regulatory network (Fig. 2E, more examples in fig. S2B). We discovered that the overall responsive interaction scores among the retinal cell types correlated with the “distress” interactions from RGCs (fig. S2, C and D). Dynamics patterns similar to distress interactions were also noted for the responsive interactions (fig. S2E). These interactions also play functional roles in axonogenesis, neuron projection guidance, and synapsis organization (fig. S2F). Together, we established a retinal cell-cell communication atlas in normal and stressed physiological conditions with temperaspatial information.

### Protective interactions associated with RGC survival

Prior studies have shown differential resilience to cell death following stress and damage in distinct transcriptome-and function-related RGC subclasses (*4, 27, 28*). Three major RGC subclasses (i.e., ipRGC, αRGC, and Gpr88RGC) demonstrate the highest survival rates post-ONC, while other RGC subclasses are more susceptible to injury resulting in cell death (*4, 23*). We sought to determine if intercellular communications between different RGC subclasses and retinal cells contribute to the distinct survival rates of RGC subtypes. By calculating the score of ligand-receptor interactions (S_LR_), established in our recent methodology study (*23*), we discovered that the overall interaction scores (sum of S_LR_) were higher for three high-survival RGC subclasses compared to low-survival RGCs (Fig. 3A). This finding suggested that in the microenvironment, neighboring retinal cells participate to regulate RGC survival in addition to previously identified cell autonomous mechanisms (*3-6*). Interestingly, the high-survival RGC subclasses expressed more ligand and receptor genes (Fig. S3A), suggesting that high-survival RGCs are intrinsically programmed to foster more communication with their surroundings.

**Fig. 3.**
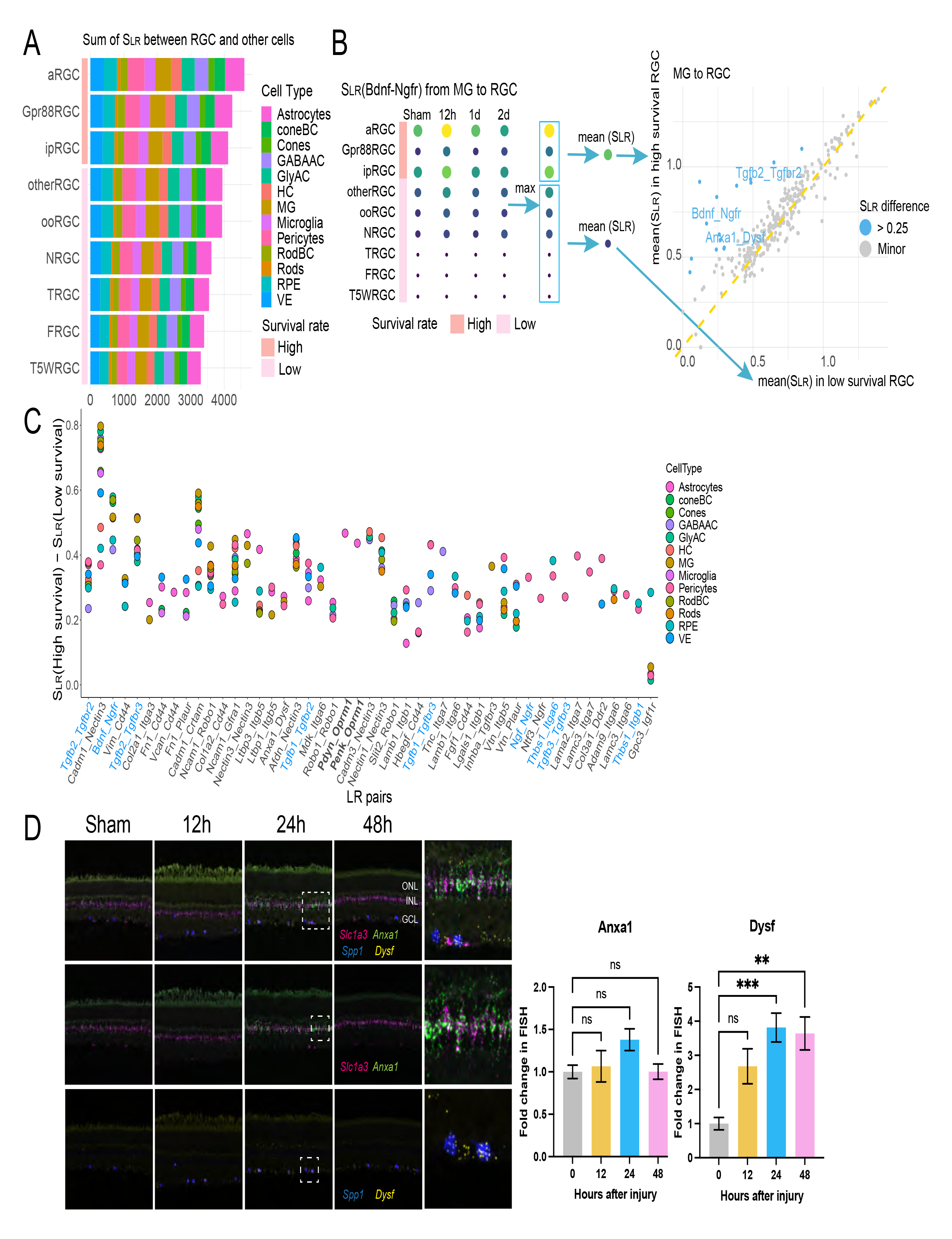
Prediction of neuroprotective interactions for RGCs. (**A**) The sum of interaction scores (S_LR_) between each RGC subclass and other retinal cell types (both from and to RGCs). The top three rows illustrate the three high-survival RGC subclasses, while the rows below them illustrate the low-survival RGC subclasses. (**B**) The workflow for calculating protective interactions sent to RGCs by Müller glia (MG), as one example cell type. Left: using the interaction *Bdnf-Ngfr* as an example, the interaction scores (S_LR_) were calculated between each RGC subclass and MG at four time points. The maximal scores across four time points (each row) were obtained for mean calculation next step. The mean values of interaction scores were then computed separately for high-and low-survival RGC subclasses (the max S_LR_ scores enclosed in two rectangles). Right: For each ligand-receptor pair, the two calculated mean values, one (y-axis) for the high-and the other (x-axis) for the low-survival RGC subclasses were plotted for comparison. Each dot represents a ligand-receptor interaction. Blue dots are the interactions that are stronger in high-than in low-survival RGC subclasses. Gray dots are the interactions with similar interaction scores between the two categories. (**C**) Summary of the potential protective interactions sent from different retinal cell types to RGCs. Similar to the calculation of MG-to-RGC protective interactions illustrated in panel (B), the neuroprotective interactions were identified in retinal cell types to RGCs. The top 47 pairs of neuroprotective interactions were identified. The known neuroprotective factors are colored blue, and the novel candidate Oprm1 receptor-related interactions are labeled font bold. (**D**) Validation of *Anxa1-Dysf* interaction, from MG to αRGC, using *in situ* hybridization. The last column of micrographs are the images with higher magnification for the boxed areas in the 24h column. The first row shows the signal of *Anxa1*, *Slc1a3, Dysf*, and *Spp1.* The second row shows the co-localization of *Anxa1* and MG marker gene *Slc1a3*, and the third row shows the co-location of *Dysf* in αRGC marker gene *Spp1*. Quantification of fluorescent intensity for *Anxa1* and *Dysf*. Data are presented as mean ± SEM. One-way ANOVA, multiple comparison, **p < 0.01, ***p < 0.001.

Next, we tried to determine the identities of individual ligand-receptor interactions sent from other retinal cell types to RGCs that might contribute to the different survival rates of RGC subclasses. To do so, we calculated the S_LR_ of all ligand-receptor interactions received by different RGC subclasses. By comparing the interaction strength (i.e., mean S_LR_ values) between high-and low-survival subclasses, we discovered some interactions were stronger (i.e., larger S_LR_) in high-survival subclasses than in low-survival subclasses, which are likely to be neuroprotective interactions (Fig. 3B). Interestingly, we did not identify any neurotoxic interactions (i.e., those with larger S_LR_ in low survival subclasses). For example, for the signals from Müller glia to RGCs, we identified 15 interactions (blue dots in Fig. 3B right panel) favored in high-survival subclasses but none in low-survival subclasses. Among these 15 ligand-receptor pairs, the roles of TGFβ- TGFβR2 and BDNF-NGFR signaling in RGC survival have been previously reported, indicating that our analysis was successful in identifying known neuroprotective interactions (*29-31*). Similar results were also observed between other retinal cell types and RGCs (fig. S3B).

By combining the neuroprotective interactions from all retinal cell types analyzed, we identified 47 non-redundant interactions that potentially promote RGC survival post-ONC (Fig. 3C and Table 1). Most of these neuroprotective interactions are initiated by ligands expressed in multiple retinal cell types (Fig. 3C). For instance, the cell adhesion molecule Cadm1, which is expressed on almost all retinal cell types included in the study, is a ligand for the receptor Nectin3, which is preferentially expressed in the αRGCs. Conversely, some neuroprotective interactions were specific to certain retinal cell types. For instance, Penk-Oprm1, Tnc-Ltga7, and Lama2-Itga7 were protective interactions to high-survival RGCs from the specific sender cells of astrocytes, GABAergic amacrine cells, and pericytes. Our predicted protective interactions include numerous known pro-survival factors, including TGF-β, BDNF, NGF, and THBS1 (*29-33*).

**Table 1.**
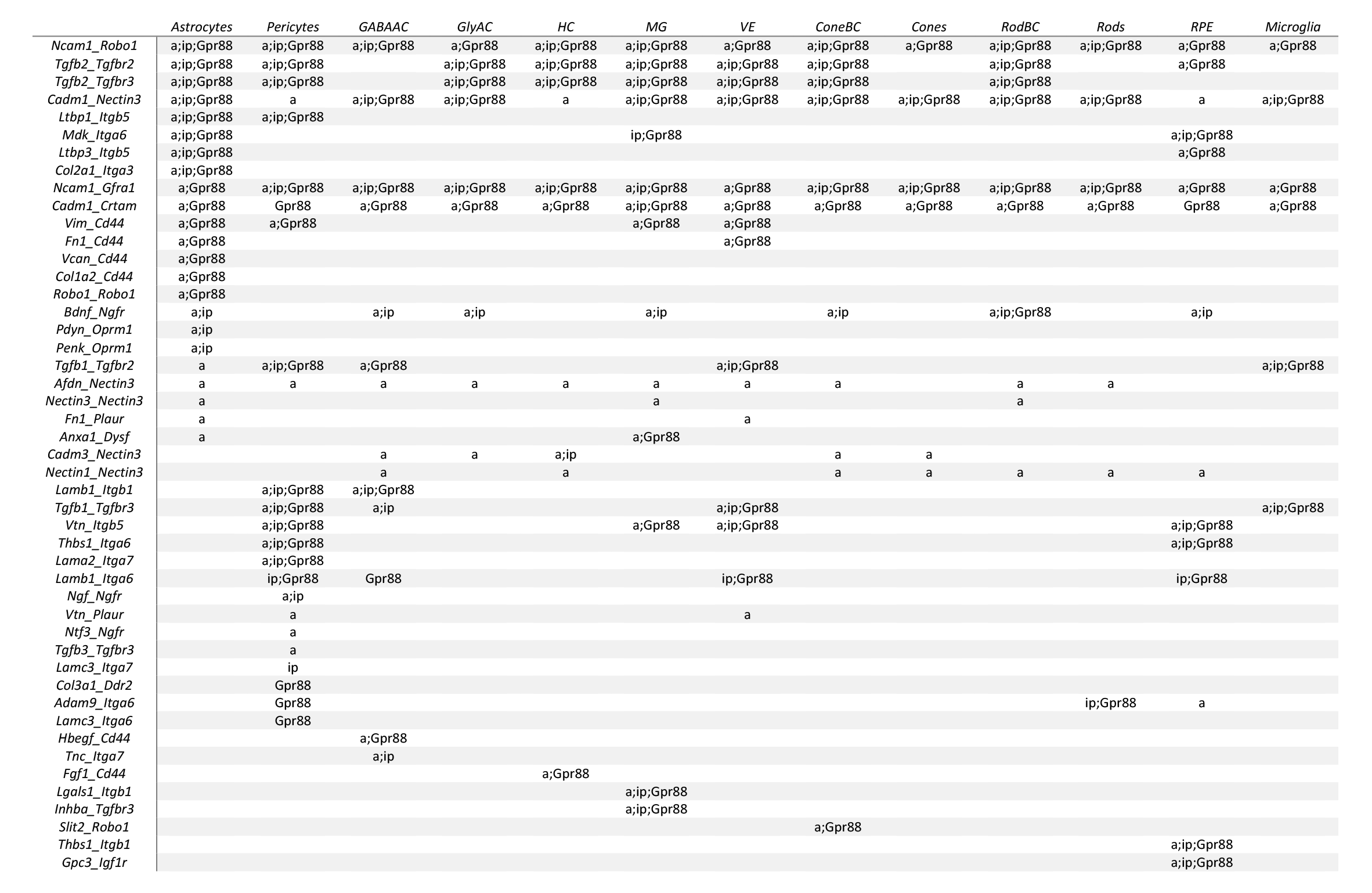

We validated the expression pattern of Anxa1-Dysf, one of the newly-identified interaction, using fluorescence *in situ* hybridization (Fig. 3D. The ligand Annexin A1 (*Anxa1*), a known regulator in inflammation (*34*), was detected in the MG, co-localizing with MG marker Slc1a3. The receptor Dysferlin (*Dysf*), which is thought to be involved in muscle regeneration and is also found to accumulate in brains of Alzheimer’s disease (*35*), was detected in αRGCs, co-localizing with Spp1 (osteopontin). One day post-ONC, a 3.9- and 1.4-fold increase of *Dysf* and *Anxa1* expression was observed at the transcriptional level compared to the sham condition, respectively (Fig. 3D). These results align with the scRNA-seq data, which showed a 2.6-fold increase for *Dysf* in αRGCs and a 1.1-fold increase for *Anxa1* in Müller glia following injury.

### Features of the protective interactions

To gain further insights into how cell-cell communications regulate RGC survival, we examined the expression patterns of ligands or receptors from these neuroprotective interactions in different RGC subclasses. We discovered that the three high-survival subclasses each had their distinct sets of receptor and ligand gene expression (Fig. 4A and fig. S4A), suggesting RGC subclasses might be intrinsically equipped to use specific ligand-receptor pairs for intercellular communications with other retinal cells. In contrast, the subclasses with low survival rates had no clear cell type specific ligand or receptor enrichment. When we examined a group of neuroprotective interactions identified with astrocytes as sender cells, we found that while some ligand-receptor paris were shared by all three high-survival RGC subclasses, the others were preferentially enriched in each RGC subclass (Fig. 4B). For instance, the Fn1-Plaur pair was stronger in αRGC, while the Fgf13- Scn5a pair showed a preference in Gpr88RGC (Fig. 4B). Our analysis suggests that high-survival RGC subclasses might use distinct cell-cell communication to promote resistance to cell death.

**Fig. 4.**
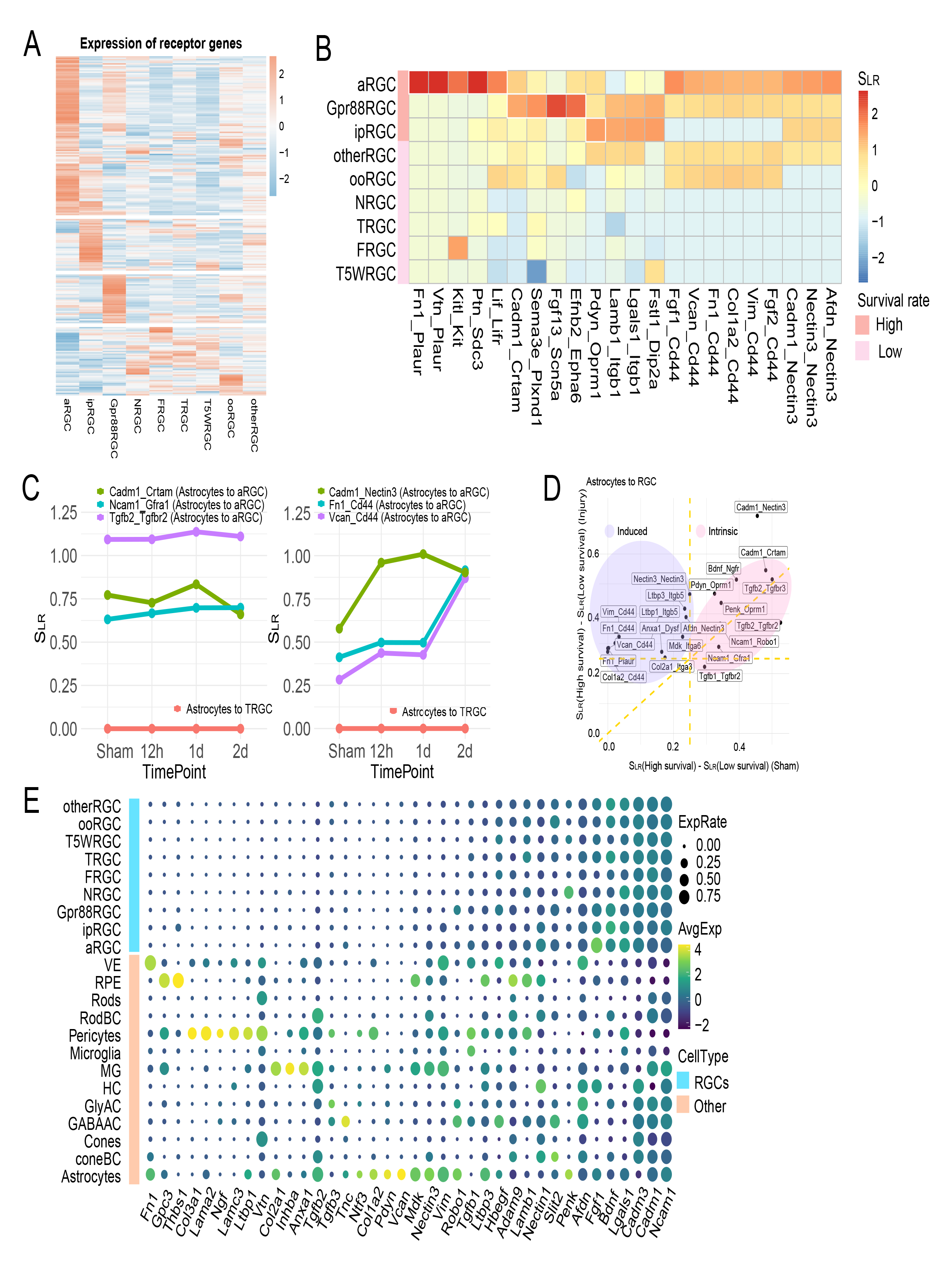
Features of the protective interactions. (**A**) Average expression level of receptor genes in RGC subclasses. (**B**) Top protective ligand-receptor pairs identified in interactions from astrocytes to RGCs. The heatmap scale represents the interaction score S_LR_. (**C**) Examples of three preset (left) and three induced (right) protective interactions. The interactions are all from astrocytes to αRGCs. Preset protective interactions are constantly higher in high-survival RGC subclasses than the low-survival subclasses, while the induced interactions become stronger after injury. Shown in pink lines, all these interactions are almost undetectable from astrocytes to T-RGC, a low-survival RGC subclass. (**D**) Summary of the preset and induced protective interactions. X-axis is the difference between interaction scores in high-and low-survival subclasses before injury; Y-axis is the difference between interaction scores after injury. (**E**) RGC autocrine *vs.* paracrine. For the ligand genes involved in protective interactions with receptors expressed on high-survival RGCs, the dot plot summarizes expression levels of ligand genes in RGC subclasses (top rows) and in other retinal cell types (bottom rows), at 12h post-ONC.

We next tested whether these protective interactions were preset in uninjured high-survival RGCs or only induced by the ONC injury. The results showed that some protective signaling interactions were active in the high-survival RGCs prior to ONC injury. For instance, Tgfb2-Tgfbr2 from astrocytes to RGCs was significantly stronger in high-survival RGCs in uninjured mice and remained largely unchanged post-injury (Fig. 4C). Some protective interactions, however, were upregulated following ONC. For example, Fn1-Cd44 signaling from astrocytes to αRGCs, was upregulated post-ONC injury, though was already stronger in uninjured high-survival RGCs. In contrast, these interactions (e.g. T-RGC) were low before and after ONC with low survival RGCs (examples shown by pink line in Fig. 4C). These protective interactions can be therefore classified into two categories based on their dynamic patterns: intrinsic and induced (Fig. 4D). Similar groups of protective ligand-recptor pairs were also observed in other retinal cell-RGC interactions, such as GABAergic amacrine cells (GABA-AC) to RGCs and MG to RGCs (fig. S4B).

Finally, we asked whether these protective interactions were primarily achieved through paracrine or autocrine signaling. To address this question, we analyzed the expression patterns of 38 ligands involved in the 47 protective interactions (Fig. 4E). Many ligands were predominantly expressed in non-RGC retinal cells, indicating that most protective interactions function through paracrine signaling. Some ligands (e.g., Fgf1, Ncam1) were expressed in both RGCs and non-RGC retinal cells, suggesting that the associated interactions could be both autocrine and paracrine. However, given the small proportion of RGCs in the total retinal cell population, these interactions could still be largely mediated by paracrine signaling.

### Oprm1 promotes RGC survival

From those top 47 ligand-receptor interactions that may contribute to RGC survival, we selected interactions involving Oprm1 for functional validation. Oprm1 is the µ-Opioid receptor, encoded by the *Oprm1* gene, preferentially expressed in intrinsically photosensitive RGCs (ipRGC), one of the high-survival subclasses. Our data showed that Oprm1 interacted with the endogenous ligands proenkephalin (Penk), prodynorphin (Pdyn), or proopiomelanocortin (Pomc), secreted by multiple retinal cell types, including astrocytes and Müller glia (see Fig. 3C).

We first examined if ectopic overexpression of *Oprm1* selectively in pan-RGCs could impact RGC survival using two established *in vivo* cell death models, ONC and the N-Methyl-D-aspartic acid (NMDA) excitotoxicity models (*3, 17, 21*). vGlut2-Cre;LSL-Sun1GFP mice were infected with either AAV2-FLEX-Sun1GFP (Sham) or AAV2-FLEX-hOprm1 for RGC specific overexpression of Oprm1. ONC was performed two weeks after the virus infection and cell survival was assessed at both 5 days and 14 days post-ONC (Fig. 5A). Oprm1 overexpression increased cell survival rate from 63.8% to 85.4%, and from 18.7% to 36.0%, at 5 and 14 days post-ONC, respectively (Fig. 5, B to D, and fig. S5A). Following NMDA injection (Fig. 5E), Oprm1 overexpression increased cell survival from 10.7% to 23.6% 7 days post-injury (Fig. 5, F and G, and fig. S5B).

**Fig. 5.**
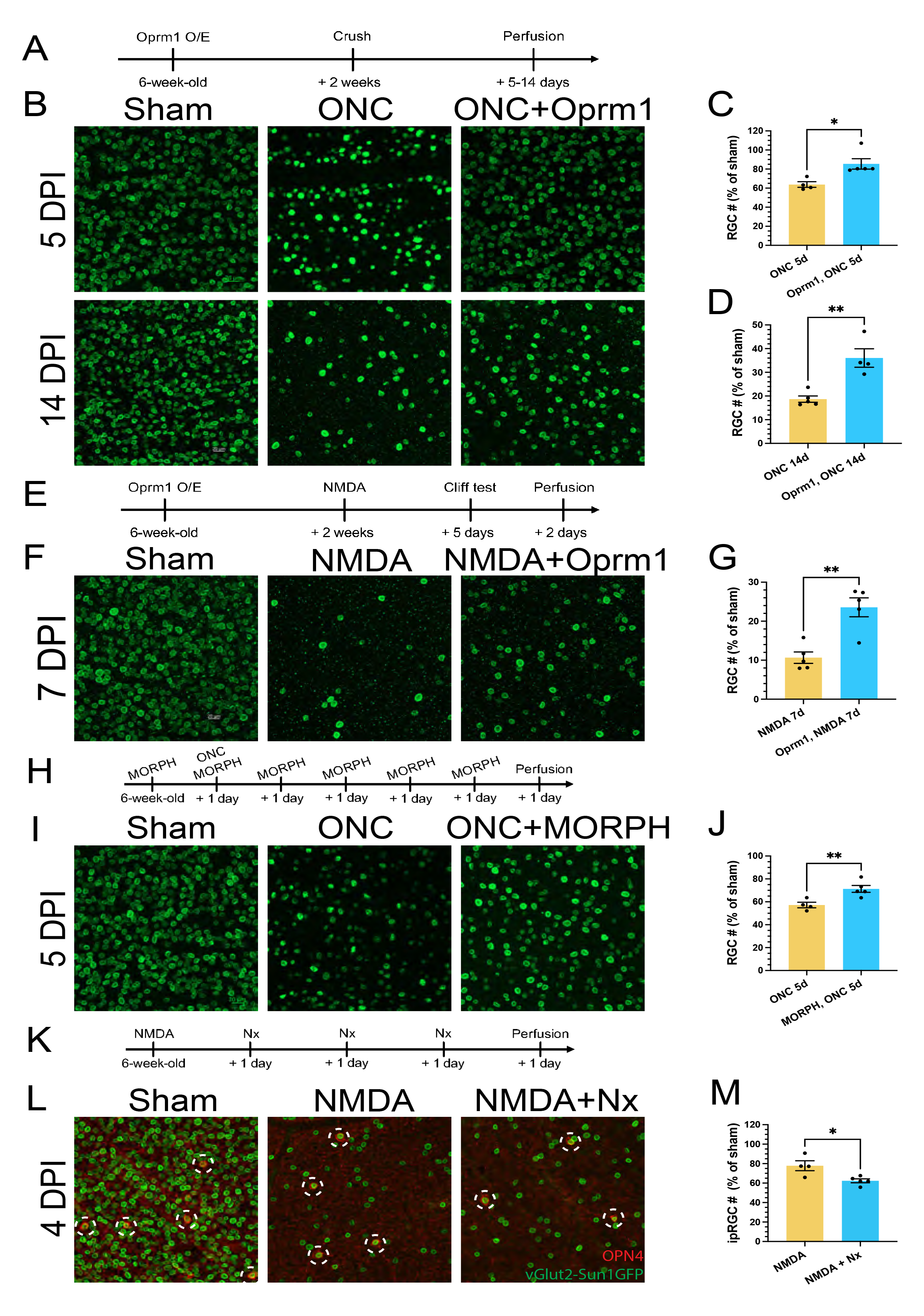
Neuroprotective effect of Oprm1 on RGC survival. (**A**) Experimental timeline of AAV2 infection, crush injury induction and sample collection in ONC model. (**B**) Oprm1 effect on RGC cell survival using ONC model. Fluorescent micrographs of pan-RGC marker vGlut2-Sun1GFP show the survived RGC cells at 5-day (upper row) or 14- day (lower row) post-ONC. (**C** and **D**) Quantification of RGC cell survival in ONC model at 5 days or 14-days post-ONC. The numbers were normalized by the RGC cell counts in sham condition. (**E**) Experimental timeline of AAV2 infection, NMDA damage, sample collection in NMDA model. (**F**) Oprm1 effect in RGC cell survival in NMDA model. (**G**) Quantification of cell survival in NMDA model. (**H**) Experimental timeline of morphine treatments, crush injury induction and sample collection. (**I**) Morphine effect on RGC cell survival at 5 days post-ONC. (**J**) Quantification of cell survival with morphine treatment. (**K**) Experimental timeline of NMDA damage, naloxone treatment, and sample collection. (**L**) Naloxone effect on ipRGC cell survival in NMDA model. Fluorescent micrographs of OPN4 and pan-RGC marker vGlut2-Sun1GFP show the survival of ipRGCs (OPN4+). The red channel represents OPN4 staining, and green represents nucleus membrane GFP in RGCs. White circles with dash line mark the ipRGCs. (**M**) Quantification of ipRGC cell survival with naloxone treatment. Data are presented in C, D, G, J, M as mean ± SEM. One-way ANOVA, multiple comparison, *p < 0.05, **p < 0.01.

We next used pharmacological reagents to confirm the neuroprotective role of Oprm1. The effects of Oprm1 agonist morphine on RGC protection after ONC were evaluated by daily intraperitoneal injection (Fig. 5H). Morphine application enhanced RGC survival from 57.2% to 71.3% 5 days post-ONC (Fig. 5, I and J). Conversely, we utilized naloxone, a potent Oprm1 antagonists, to test if the Oprm1 activation was necessary for RGC survival post-ONC. Since *Oprm1* is preferentially expressed in high survival ipRGCs, we focused on determineing how naloxone affected ipRGC survival following NMDA excitotoxity. As anticipated, the ipRGC survival rate further decreased from 77.8% to 62.4% following daily intraperitoneal naloxone injection (Fig. 5, K to M and fig. S5C). Similar reductions in ipRGC survival were also observed following ONC (fig. S5, D to F). Collectively, overexpression of Oprm1 had neuroprotective effects on RGCs after 2 different types of retinal injuries.

### Universal protective effect of Oprm1 on different RGC subclasses

To gain insights into molecular mechanisms by which Oprm1 protects against ONC, we performed the single-nucleus RNA sequencing (snRNA-seq) from enriched RGC nuclei (vGlut2-Cre;LSL-Sun1GFP) in the ONC model. We collected three samples from sham condition, ONC model (5 days), and RGC overexpressing *Oprm1* in ONC model (5 days). The RGC nuclei enrichment was successful (fig. S6, A and B). In these samples, a total of 6348, 3314, and 6141, respectively, RGC nuclei were profiled, representing all major RGC subclasses (Fig. 6A). In line with our intercellular communication analysis, we observed that the endogenous *Oprm1* was predominantly expressed in ipRGCs and a subset of αRGC (Fig. 6, B and C), while the virus-transduced overexpression of human *Oprm1* was detected in all subclasses of RGCs (Fig. 6B and fig. S6C).

**Fig. 6.**
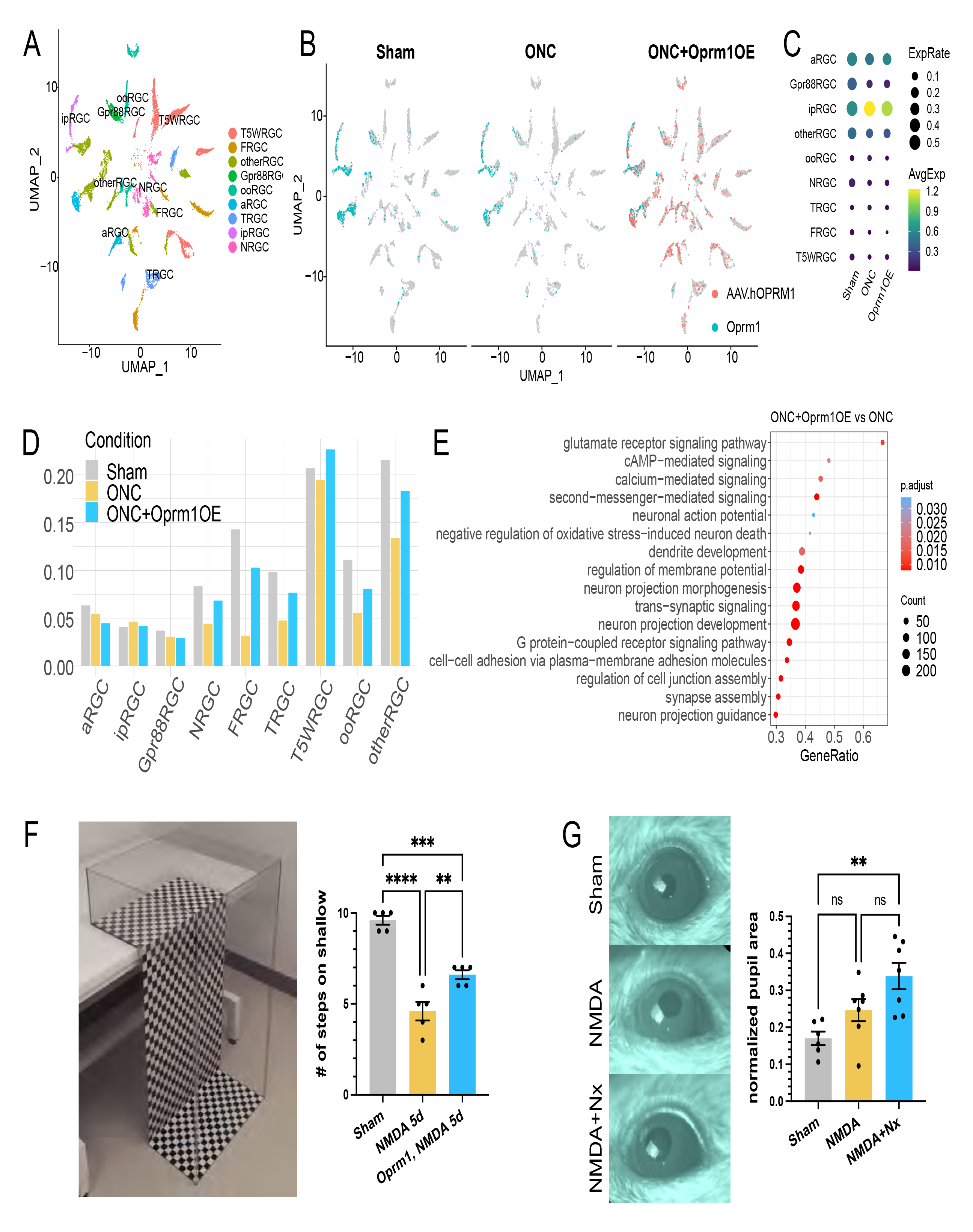
snRNA-seq of RGCs after Oprm1 overexpression. (**A**) Clustering of RGCs showing the RGC subclasses. Samples collected from sham, 5-day post-ONC, and ONC with Oprm1 overexpression conditions were integrated. (**B**) Expression patterns of endogenous Oprm1 (cyan) and/or sequenced AAV WPRE element standing for the ectopic expression of human Oprm1 (pink) in RGC subclasses under each condition. (**C**) The dot plot for expression level of endogenous Oprm1 in RGC subclasses in three conditions. (**D**) Normalized cell type proportions among RGC subclasses in three conditions. (**E**) Representative biological processes enriched in the Oprm1 overexpression samples. (**F**) Visual cliff behavioral test on the effect of Oprm1 overexpression. Left: the equipment setup of visual cliff test. Right: the average occurrences of mice choosing the shallow side (indicator of acute vision) out of 10 trials. Each dot represents one mouse. One-Way ANOVA, multiple comparison **p < 0.01, ***p < 0.001, ****p < 0.0001. (**G**) Pupil reflex test on naloxone effect in NMDA model. Left: representative images of pupil in three conditions. Right: normalized pupil size in three conditions. One-Way ANOVA, multiple comparison, **p < 0.01

We next investigated which RGC subclasses were protected from ONC by *Oprm1* overexpression. To normalize cell counts across each condition, we first computed the cell percentages in each RGC subclass and then adjusted these percentages in the control ONC and *Oprm1*-overexpressed ONC samples by 63.8% and 85.4%, respectively, based on total survival rates determined by immunostaining (see Fig. 5C). Because most cells within the high-survival subclasses (ipRGC, αRGC, and Gpr88RGC) were alive 5 days post-ONC, we did not observe neuroprotective effect additionally by Oprm1 overexpression on these subclasses (Fig. 6D). For low survival subclasses, the cell numbers significantly declined following ONC, but *Oprm1* overexpression substantially reduced cell death, indicating a broad protective effect of Oprm1 on all RGC subclasses (Fig. 6D).

Finally, we analyzed the downstream pathways activated by *Oprm1* overexpression. By comparing the gene expression between ONC and *Oprm1*-overexpressed ONC samples, we identified 337 differentially expressed genes. Gene Ontology (GO) analysis revealed that the biological processes enriched in *Oprm1*-overexpressed sample were related to cAMP and/or calcium second messenger signaling, negative regulation of neuron death, regulations of synaptic transmission and membrane potential, as well as neuron projection morphogenesis and guidance (Fig. 6E).

### Oprm1 maintains visual functions following RGC injury

We then investigated if neuroprotection induced by Oprm1 improved visual function. First, we used the visual cliff test, which evaluates visual acuity based on the mouse’s innate response to avoid the visually deep area. Since the ONC completely disrupts RGC projections to the brain, the visual cliff test was employed specifically with the NMDA model that did not completely damage the visual pathway. Mice were placed at the edge, or “cliff”, between shallow and deep sides, and their preference for either side were recorded. We examined groups of five mice, with each mouse undergoing ten trials. Uninjured mice, on average, chose the shallow side 9.6 times out of 10 trials, in sharp contrast to the NMDA-treated group’s average of 4.6 times (Fig. 6F). However, Oprm1 overexpression significantly improved the shallow side selection to 6.6 times on average in the NMDA-treated mice (Fig. 6F).

ipRGCs are the major relay for light signals to reach brain centers that regulate the pupillary light response (PLR), and therefore we used the PLR to examine whether naloxone-induced ipRGC loss showed a physiological outcome on ipRGC signaling in mice. We employed sustained PLR as a marker to evaluate the status of ipRGCs in NMDA-injured retinas (*36*). Althoug a decrease in the total number of ipRGC cells was observed after an intravitreal NMDA injection (Fig. 5, K to M), the resultant ipRGC numbers were not sufficient to induce PLR deficits (Fig. 6G), corroborating previous observations from Opn4aDTA mice with partial loss of ipRGC (*37*). However, when naloxone treatment was combined with NMDA, more substantial reduction in ipRGC numbers were observed (Fig. 5, K to M), and PLR deficits were detected (p=0.001, n=7), implying that the Oprm1-mediated ipRGC protection is attenuated by anti-opioid activity at the functional level.

### Protective effects of Oprm1 in a glaucoma model

Finally, we investigated if Oprm1 overexpression in RGCs protected neurons in an experimental model of pressure-induced glaucoma. The previously established magnetic microbeads occlusion model (*38*) was employed to increase the intraocular pressure (IOP). Specifcially, the beads were intracamerally injected into the anterior chamber, adhering between the circumference of cornea and iris, and obstructing the circulation of aqueous humor (Fig. 7A). An elevation in IOP was then maintained for a period of eight weeks (*3, 38*) (Fig. 7B). Similar to the previous cell death models, AAV2-mediated Oprm1 overexpression specifically in pan-RGCs enhanced RGC survival in this glaucoma model at eight weeks after IOP elevation. While the survival rate of RGC in the elevated IOP (eIOP) group was 62.2%, Oprm1 overexpression in RGCs significantly enhanced the survival rate to 84.0% (Fig. 7, C and D, and fig. S7, A and B).

**Fig. 7.**
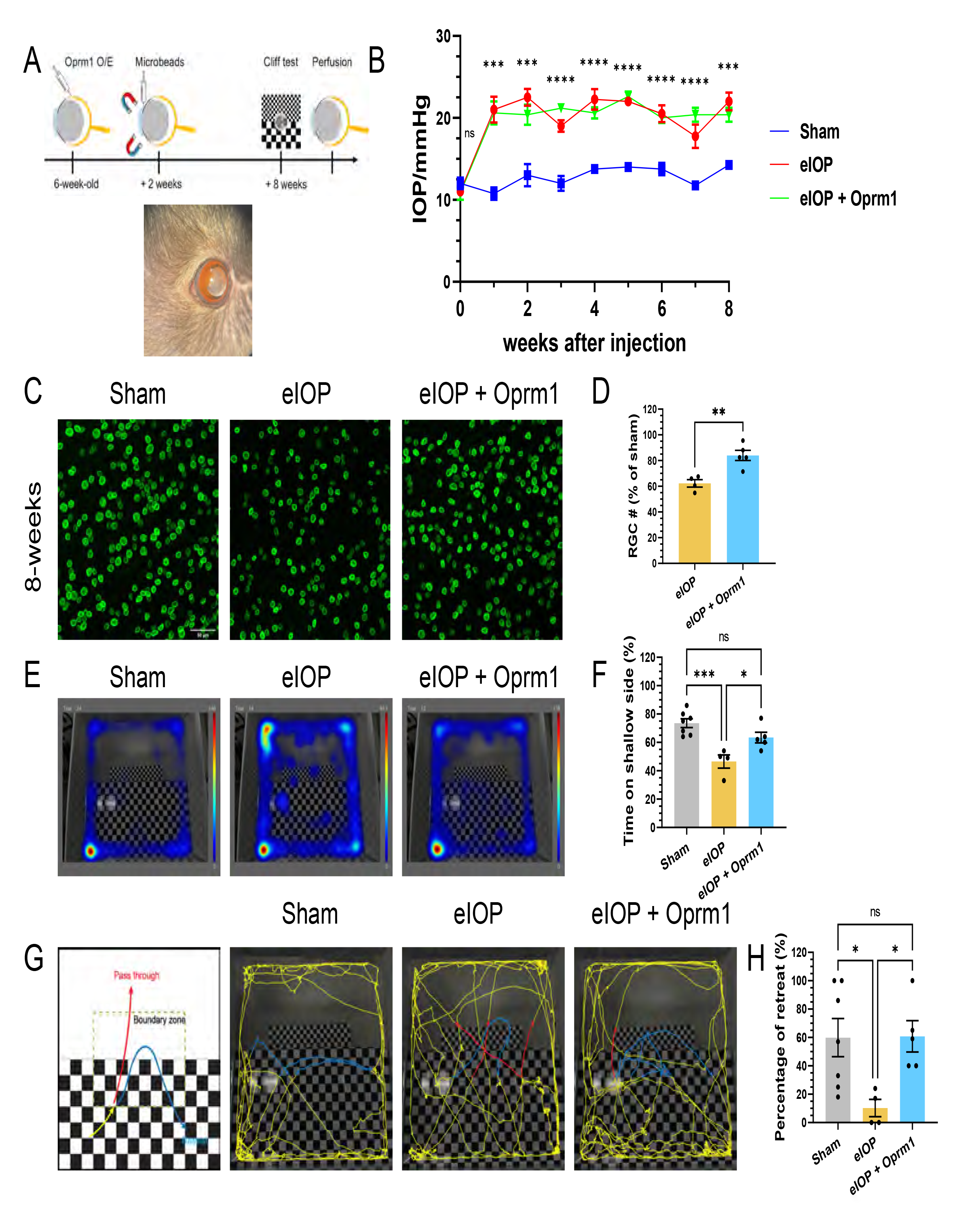
Oprm1 effect in a glaucoma model. (**A**) Experimental model for elevation of intraocular pressure (IOP) induced by intracameral injection of magnetic microbeads. A representative photo shows magnetic microbeads stuck between the circumference of cornea and iris after injection. (**B**) IOP in 8 weeks. Data are presented as mean ± SEM. Sham group, n=7; eIOP group, n=4; Oprm1 + eIOP, n=5. One-way ANOVA, ***p < 0.001, ****p < 0.0001. (**C** and **D**) Protection of RGCs from eIOP with overexpression of Oprm1. Scale bar, 50µm. (**E-H**) Assessment of Oprm1 overexpression on visual function using cliff test. (**E**) Representative movement heatmaps from sham, eIOP and Oprm1 + eIOP groups. (**F**) Quantification of time spent on the shallow side during the 30-minute period in the test. (**G**) A schematic diagram and representative traces (5 min) of the visual cliff test. Red traces highlight “pass through” from the shallow side to the deep side, while blue traces highlight “retreat” to the shallow side. (**H**) Percentages of retreat events among all traces entering the boundary zone during the 30-minute period. Data are presented as mean ± SEM. Sham group, n=7; eIOP group, n=4; Oprm1 + eIOP, n=5. One-Way ANOVA, multiple comparison, *p < 0.05, **p < 0.01.

To assess how visual functions in the glaucoma model was affected by Oprm1 overexpression at eight weeks following microbeads injection, we used two different visual-based behavioral assays. We first tracked the free movement of mice in the visual cliff test arena over a 30-minute period, and analyzed their location distribution throughout this time. The uninjured sham group spent 73.6% of the time on the shallow side of the arena, whereas the mice in eIOP group spent only 46.5% of the time there, indicating reduced depth perception. When Oprm1 was overexpressed in eIOP mice, the time on the shallow side was significantly increased to 63.4% (Fig. 7, E and F). Second, we examined if the mice would turn away from or pass through the shallow-to-deep “cliff” moving from the shallow to the deep side (Fig. 7G). The results showed that when entering the cliff zone from the shallow side, uninjured mice had 59.9% chance to retreat, while the eIOP injured mice merely had 10.3% chance to retreat. However, the eIOP mice with overexpressed Oprm1 had 60.8% chance to retreat, comparable to that of uninjured mice (Fig. 7H). The results indicates that Oprm1 overexpression in RGCs improves the vision acuity in the visual cliff test, likely through its role in RGC neuroprotection.

## DISCUSSION

### Organizing principles of cell-cell communication

In this work, we established a systematic cell-cell communication atlas between RGCs and other retinal cell types in response to retinal injury. Our analysis has identified spatiotemporal organizing principles for cell-cell communications within a complex system composed of diverse cell types. First, we revealed that the numbers of ligand-receptor interactions between RGCs and other cell types varied based on the relative positions within the retina. Specifically, Müller glial cells and the astrocytes, which contact directly with RGCs, have the highest level of communication. In contrast, cell types that were physically separated from RGCs, such as cones and rods, have far fewer interactions. Second, we identified temporal variation in cell-cell communication following optic nerve crush. Even though some cell-cell interactions were transient, returning to their original states quickly, we anticipate that the retinal cells received the distress signals from RGCs, and the response was already trigged through the short period of deviation from the original states.

### Cell-cell interactions as intrinsic cellular property

While the microenvironment was extensively studied in the cancer field, stem cell biology, and immunology, it has not been systematically analyzed in a mature nervous system. In this work, we identified the neuroprotective factors from the perspective of analyzing cell-cell communication. One interesting finding is that the high-survival RGCs have more intercellular interactions than those with low survival abilities after injury. Furthermore, high-survival RGC subclasses express more ligands and receptors, suggesting that the property of cellular resilience to cell death is, in part, encoded within the cell-cell communication and established during development. Indeed, the concept of “intrinsic” cellular properties are not constricted within insular cells. In a broad sense, the interactions with neighboring cells are actually another aspect of the intrinsic cellular property.

### All neighbors are good neighbors

In our analysis, we identified neuroprotective interactions from retinal cells to RGCs by comparing the interaction strengths between high-and low-survival RGC subclasses. It was unexpected that we only found retinal cell-to-RGC interactions that were stronger in high-survival RGCs than in low-survival RGCs. We did not find any preferential interactions that were in low-survival RGCs, which likely contribute to some neurotoxic effects for the low-survival RGCs. In other words, the neighboring retinal cells are helping to promote RGC survival post-injury, rather than sending any additional apoptotic signals to destroy injured RGCs. This is a stark contrast to the transcriptome comparison among RGC subclasses, in which the crush operation induces differentially expressed genes significantly in either high-or low-survival subclasses of RGCs.

### An old receptor with a new function

We functionally validated one of the predicted neuroprotective factors identified in this analysis - the μ-opioid receptor encoded by *Oprm1* gene. Despite its well-documented role in pain regulation, reward, and addictive behaviors (*18-20*), Oprm1 has not been previously implicated in regulating neuronal survival. Both gain-and loss-of-function studies demonstrated that Oprm1 is sufficient and necessary to promote and support RGC survival following multiple types of injury models. Furthermore, although Oprm1 is predominantly expressed in intrinsically photosensitive RGCs (ipRGCs), overexpression in other RGCs can also protect more vulnerable RGC subtypes. Notably, Oprm1 overexpression in RGCs significantly improved visual perception in a high intraocular pressure-induced glaucoma model, suggesting its future translational potential. The high-affinity selective agonists for Oprm1 that are already in clinical use for pain management may therefore be potentially repurposed for RGC neuroprotection in emergency situations following acute optic nerve injury or in closed-angle glaucoma. In addition, Oprm1 is highly expressed in ipRGCs, exerting its neuroprotective role after various type of retinal injuries. It is very likely that activation of Oprm1 as a potential therapeutic approach would have lower side effects.

Together, our study established an effective strategy to discover functionally important cell-cell communication in a complex *in vivo* system. Moreover, the combined experimental and analytic platform will facilitate the mechanistic study of intercellular interactions in diverse biological systems beyond the nervous system.

## METHODS

### Mice

All animal experiments were approved by the Institutional Animal Care and Use Committees (IACUC) at Johns Hopkins University School of Medicine. The mice were maintained in a climate and light/dark (14h/10h) controlled facility, with continuous access to food and water. All studies were conducted on adult mice 2- to 3- month age and on both male and female mice. The following mouse strains were utilized - C57BL/6 mice (JAX #000664), vGlut2-IRES-Cre (JAX #016963), and CAG-LSL-Sun1/sfGFP mice (JAX #021039), all available at the Jackson Laboratory (Bar Harbor, ME). Homozygotes vGlut2-Cre;LSL-Sun1GFP were generated by breeding and backcross.

### Optic nerve crush (ONC)

After anesthesia with Avertin (200-500 mg/kg intraperitoneal injection), the right optic nerve was exposed intraorbitally and crushed with #5 Dumont forceps (FST, Foster City, CA) gently at ∼1mm behind the eyeball. The sham operation was conducted with the same anesthesia procedure and optic nerve exposure under the microscope, without crushing of the optic nerve. All microsurgeries were performed by experienced personnel using the Leica M80 stereo microscope.

### Retinal tissue dissociation and Droplet-based scRNA-seq

With 12hour-, 1day-, 2day-ONC, or sham conditions, the mice were euthanized. Each time point had two biological replicates of mice. The eyeballs were enucleated and kept in ice-cold PBS. The retinas were quickly and carefully dissected, and whole retinal cells were dissociated using the Papain enzyme system (LK003150, Worthington). 30μM actinomycin-D was added to the Papain system to prevent post-enucleation transcription. The tissues then underwent manual trituration in neurobasal medium supplemented with 3% BSA, 10 mg/mL Ovomucoid, 1U/uL RNase inhibitor, and 3μM actinomycin D. Cells were filtered through a 40-μm strainer and centrifuged under 500×g. For scRNA-seq, the cell concentration was kept at ∼1000 cells/uL. The scRNA-seq libraries were prepared using the 10X Genomics Chromium 3’ Single-Cell Gene Expression V3 Kit following the standard protocol by the manufacturer. Approximately 8,000-12,000 cells were loaded onto the platform. Libraries were sequenced with the NovaSeq S2 100 platform.

### Quality control, clustering, and cell type identification of scRNA-seq data

Raw reads of adult mouse retina scRNA-seq data before and post-ONC (control, 12h, 1d, and 2d) were mapped to mm10 and expression matrices were generated using Cell Ranger 3.0.2 from 10X Genomics. In addition, we downloaded the corresponding RGC data from GEO (GSE137400). We created a Seurat object for each sample using Seurat v4.0.5. For the whole retina dataset, cells were removed if their nUMI was <800 or >30000, nGene was <350 or >7500, mitochondrial gene rate was >20%, or log10GenesPerUMI was <0.8. For the RGC dataset, we removed cells with nUMI <500, nGene <250, mitochondrial rate >20%, or log10GenesPerUMI <0.8. We then used Scrublet v0.2.3 to remove predicted doublets with default parameters for both data sets. Furthermore, genes detected in less than 5 cells in each sample were also removed from the data. As a result, 56,531 cells from the whole retina dataset and 86,426 cells from the RGC dataset were obtained for downstream analysis. For each dataset, Seurat objects of the samples were then normalized, integrated, clustered, and visualized through UMAP using Seurat v4.0.5. Cell identities were then assigned based on the known canonical marker genes of retinal cell types.

### Magnetic purification of RGC nuclei for Droplet-based snRNA-seq

The vGlut2-Cre;LSL-Sun1GFP mice were used for this experiment. Mice under sham condition, post-ONC (5d) and post-ONC (5d) with hOprm1 overexpression conditions were euthanized. The retinas were quickly and carefully dissected out from enucleated eyeballs. Through gentle pipette trituration, the retinas were dissociated in ice-cold neurobasal medium on ice. According to the 10X Genomics protocol, whole retinal cells were then incubated in the nuclear lysis buffer on ice for 3-min to expose and penetrate the cell nuclei, and then centrifuged to pellet the nuclei. The cell nuclei were then resuspended in ice-cold MACS buffer (Miltenyi Biotec, Gaithersburg, MD) for anti-GFP MACS microbeads incubation. The whole retina nuclei were then passed through and eluted from the Miltenyi MACS MS column to enrich GFP+ RGC nuclei, then being centrifuged to pellet. The pellet was then resuspended to reach concentration of 4000∼5000 nuclei/uL loading onto the 10X Genomics Single-Nuclei 3’ HT platform.

### Data analysis of snRNA-seq of purified RGCs

For enriched RGC snRNA-seq data from sham condition, post-ONC (5d), and post-ONC (5d) with hOprm1 overexpression samples, raw reads were mapped to mm10, and expression matrices were generated using Cell Ranger 7.0.0 from 10x Genomics. A Seurat object for each sample was created using Seurat v4.3.0. For sham data, cells were removed if nUMI was < 1000 or > 50000, nGene was < 500 or > 10000, mitochondrial rate was > 10%, or log10GenesPerUMI was < 0.8. For ONC and ONC with *Oprm1* overexpression data, cells were removed if nUMI was < 1000 or > 30000, nGene was < 500 or > 7000, mitochondrial rate was > 10%, or log10GenesPerUMI was < 0.8. We then removed predicted doublets using DoubletFinder v2.0.3 with its standard pipeline for each sample (*39*). In total, 26170 cells were used for downstream analysis. Specifically, 9306, 6060, and 10804 cells were obtained from the sham, ONC, and ONC with Oprm1 overexpression samples, respectively. The filtered Seurat objects were then normalized, integrated, clustered, and visualized through UMAP using Seurat v4.3.0. Cell type identities were assigned based on the known canonical marker genes of retinal cell types. After cell type annotation, 15803 RGCs were obtained, with 6348, 3314, and 6141 RGCs from the sham, ONC, and ONC with Oprm1 overexpression samples, respectively. We then took the subset of the Seurat objects of each sample by removing non-RGCs and performed normalization, integration, clustering, and UMAP visualization again using Seurat. RGC subtype identities were then assigned based on the projection of the RGC atlas data from Tran et al. (*4*) onto our clusters using the Seurat data transfer pipeline and known subtype marker genes.

### Prediction of ligand-receptor interactions between RGCs and other cells

We predicted ligand-receptor interactions between RGCs and other retinal cells, as well as between each RGC subclass and other cells, using the package LRLoop, with detailed algorithm in (*23*). In essence, the LRLoop method mainly relied on three types of information to predict ligand-receptor interactions: 1) The expression levels of ligands in sender cells and receptors in receiver cells; 2) The cell type-specific signaling networks and regulatory networks by superimposing the known networks and the expression levels of the genes in signaling and regulatory networks in a particular cell type; 3) The ligand-receptor interactions form some loops (Fig. 2E, and fig. S2B) through the signaling and regulatory networks. We first derived the interaction strength of each ligand-receptor interaction using the first two pieces of information. We then modified the strength by considering the other ligand-receptor interactions that can form a loop with the interaction of interest. Finally, the candidate ligand-receptor interactions are further filtered based on the following criteria: 1) both the ligand and the receptor genes are detected in at least 10% of the corresponding sender and receiver cell types, respectively; 2) The interaction score calculated from LRLoop is no less than 0.5 in any time point. The overall score between each pair of cell types was defined as the sum of interaction scores of identified ligand-receptor pairs.

### Differential ligand-receptor interactions between high-and low-survival RGC subclasses

For the predicted ligand-receptor interactions from each non-RGC retinal cell-type ct_1_ to RGC subclasses, DS_LR_, the interaction score difference of each LR pair between high and low survival RGC subclasses was defined as:

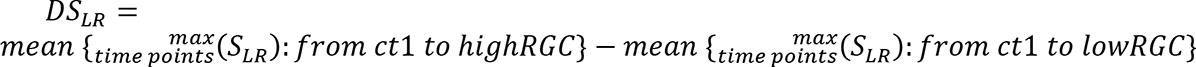

Interactions with score difference above or below cutoffs (0.25 and -0.25) were considered as differential ligand-receptor interactions between high-and low-survival RGC subclasses.

### Fluorescence *in situ* hybridization (FISH)

Eyes with optic nerves were surgically removed from perfused mice, and retinas were dissected out and post-fixed in 4% PFA at 4°C overnight. After a serial transfer in 10%, 20%, and 30% sucrose solution, dehydrated retinas were embedded in O.C.T. compound (REF 4583, Sakura) for cryosectioning into 10μm cross-sections. Fluorescent *in situ* hybridization (FISH) was performed using the commercially available RNAscope Fluorescent Multiplex assay (ACDbio) according to the manufacturer’s instruction. In brief, the O.C.T. compound was removed by washing with 1×PBS, followed by baking the slides for 30 minutes at 60°C. Sample slides were then post-fixed with 4% PFA for 15 minutes at 4°C, and serially dehydrated with 50%, 70%, and 100% ethanol for 5 minutes at room temperature. Samples were covered with hydrogen peroxide for 10 minutes, followed by boiling in 1× Target Retrieval Reagents for 5 minutes. The samples were incubated with Protease III of the HybEZ^TM^ II Hybridization System for 30 minutes at 40°C. After incubation with RNAscope probes for 2 hours at 40°C and subsequent AMP, the samples were applied with HRP, Opal fluorescent dyes, and HRP Blocker for corresponding channels, sequentially. The sample slides were mounted with Fluoroshield and coverslips. Confocal images of the slides were acquired using a Zeiss LSM 880 microscope.

### Quantification of fluorescence *in situ* hybridization

Confocal images were acquired using a Zeiss LSM 880 microscope (JHU Institute for Basic Biomedical Sciences Microscope Facility) under the same parameter settings. In addition to DAPI, four microscope channels were used: Fluorescein for Opal520 (FP1487001KT, Akoya Biosciences) to show ligand expression (Manual Assay RNAscope Multiplex Probe for Anxa1, 509291-C1, ACD Bio), Cyanine 3 for Opal570 (FP1488001KT) to show receptor expression (Dysf, 1134891-C2), Texas Red for Opal620 (FP1495001KT) to show Müller glia with a probe against Slc1a3 (430781-C3), Cyanine 5.5 for Opal690 (FP1497001KT) to show αRGCs with a probe against Spp1 (435191-C4).

Quantification of fluorescence *in situ* hybridization was obtained based on the guidelines provided by the RNAscope Fluorescent Multiplex assay (ACDbio). In brief, a threshold was set to define the shape of each discrete signal dot after the average background intensity was calculated. The average intensity per single dot was calculated within a representative region containing around 20 signal dots by the following equation:

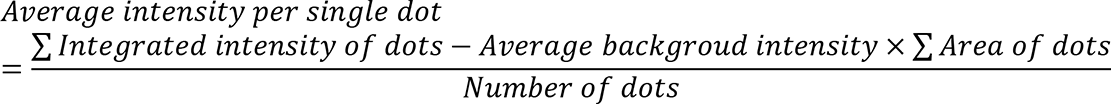

Then the total dot number in a region above threshold was calculated by the following equation:

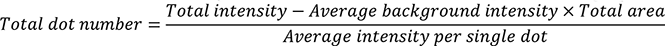

### Intravitreal injection of AAV, NMDA, and intraperitoneal injection of morphine

The intravitreal viral injection was performed as previously described (*40*). Under anesthesia, 1.5 μL of AAV2 virus was injected into the right vitreous humor of a mouse with a Hamilton syringe assembled with a 32-gauge needle. The position and direction of the injection were well-controlled to avoid injury to the lens. AAV2-FLEX-hOprm1 (Addgene plasmid# 166970) virus or AAV2- FLEX-sun1GFP (Addgene plasmid# 160141) control virus were packaged by SignaGen at high-titer (1 × 10^3^ gc/mL) to produce AAV2 viruses. For the NMDA excitotoxicity model, the mice were intravitreally injected with 1.5 μl of 20 mM N-methyl-d-aspartic acid (NMDA) (Sigma-Aldrich, St. Louis, MO). The 1.0mg/mL commercially available morphine solution (M-005-1ML in methanol, Sigma Aldrich, MO) was diluted in PBS for intraperitoneal injection into mice to reach 0.5mg/kg final concentration.

### Immunohistochemistry and microscopy

Eyes with optic nerve were surgically removed from perfused mice, followed by post-fixation in 4% PFA at 4°C overnight. Retinas were dissected out in 1×PBS and cut with four radial incisions to create a petal shape. Cold (-20°C) methanol was used to fix and flatten the retinas. Retinas were transferred into a 24-wells plate with cold methanol for storage (*41*). For immunohistochemistry, retinas were blocked in blocking solution containing 0.3% Triton X-100, 0.2% BSA and 5% goat serum in PBS at room temperature for 1 hour, followed by incubation of primary antibodies at 4°C overnight. After washing 4 × 10 minutes in PBS with 0.3% Triton X-100, retinas were incubated with secondary antibodies at room temperature for 4 hours. Finally, retinas were washed 4 × 15 minutes in PBS with 0.3% Triton X-100 and mounted with Fluoroshield (F6182, Sigma Aldrich). Confocal images were acquired using a Zeiss LSM 800 microscope (JHU neuroscience MPI core). For RGC numbers counting, eight squares (320μm × 320μm) were sampled evenly around the peripheral region of each retina. The mean cell number out of these eight positions was viewed as RGC cell number of one retina as one biological replicate.

### Elevation of intraocular pressure (IOP) by intracameral injection of microbeads

Elevation of IOP was induced by intracameral injection of magnetic microbeads into the right eye of the VGlut2-Cre; LSL-Sun1-sfGFP mice. In brief, superparamagnetic polystyrene beads (4.5μm, Dynabeads^®^ M-450 Epoxy, Invitrogen, 14011) were treated with 0.02M sodium hydroxide (NaOH) in 10× Tris buffer to remove epoxy groups. Then, the microbeads were homogenized in a sterile balanced salt solution at a concentration of 1.6× 10^6^ beads/μL. Three-month-old mice were anesthetized with an intraperitoneal injection of proper volume of Avertin (2,2,2-Tribromoethanol, T48402, Sigma Aldrich, 20mg/mL, 250mg/kg). After pupil dilation with a 1% tropicamide ophthalmic solution (NDC 70069-121-01, Somerset Therapeutics LLC, Hollywood, FL), a 30G × ½ needle (REF 305106, BD PrecisionGlide^TM^) was used to incise the edge of the cornea, and 1.5 μL of microbeads solution was injected into the anterior chamber using a sharp glass micropipette connected to a Hamilton syringe (Cat # 20919, World Precision Instruments). A small piece of the magnetic ring was applied to evenly distribute microbeads evenly around the circumference between the cornea and iris (*38*).

### IOP measurement

The IOP of both eyes was measured by the TonoLab tonometer (Colonial Medical Supply) every week. The tip of the probe was applied 1 to 4 mm from the center of the cornea after the mice were anesthetized. The IOP on each eye was measured six times, and the tonometer produced an average value of IOP automatically. All IOPs were measured around 4:00 to 5:00 pm to reduce variation.

### Visual cliff test

Mice were put into a transparent plexiglass arena (Conduct Science, Boston, MA) with a dimensions of 62cm × 62cm × 19cm (L × W × H). The arena was evenly separated into two zones: a shallow zone with a checkered pattern under it, and a deep zone with the same checkered pattern 60cm underneath it to create an illusion of depth (Figure 5F, left). NMDA-treated mice were placed in the middle line, and their choices toward different zones were recorded. The arena was cleaned between tests. For the elevated IOP glaucoma model, mice did not show significant differences in zone choices between groups two weeks after IOP elevation. Eight weeks after IOP elevation, the mice tended to stay at the starting point without much movement when placed into the arena, which may have been caused by the weekly anesthesia with Isoflurane for IOP measurement. Such behavior is consistent with previous reports showing that repetitive applications of Isoflurane could lead to cognition impairment (*42, 43*). For these mice, 30-minute videos of their movement were recorded after a 5-minute adaptation time. EthoVision XT 17 (Noduls Information Technology Inc.) was used to record and analyze videos.

Trace analysis was performed based on the videos recording mice locomotion. Two types of behavior were analyzed. First, we calculated the time the mice stayed in either the shallow or the deep zone. We then analyzed the movement in the boundary zone located in the center of the arena because the mice tended to move in a circulating way along the borders, but showed different behaviors when in the center of the arena. The boundary zone was defined as a rectangle extending 3 grids into both the shallow and deep sides from the middle line, and 3 away from the lateral borders of the area (the length of 3 grids is nearly the average body length of the mice without tails, Figure 7). The traces of mice entering the boundary zone from the shallow side were counted. If mice leave the boundary zone to the deep side, the trace is counted as a “pass through.” If mice leave from the zone to the shallow side, it is counted as a “retreat”. The ratio of retreats to total traces was calculated.

### Intraperitoneal injection of Naloxone and ipRGC number analysis

Six-week-old vGlut2-Cre; Sun1-GFP mice were used in experiments related to naloxone. For the NMDA damage model, 0.4mL of 0.5mg/mL Naloxone (0.2mg/mouse) (*44, 45*) was intraperitoneally injected daily, and mice were sacrificed four days after NMDA damage. For the ONC model, 0.4mL of 0.5mg/mL Naloxone was intraperitoneally injected daily, and mice were sacrificed five days after injury. Opn4 antibodies (PA1-780, Invitrogen) were used to label ipRGCs in immunostaining. Photos of whole mounted retina were token with a Zeiss LSM 880 Confocal Microscope. Numbers of ipRGC in each retina were counted.

### Pupillometry

Mice for pupillometry were 8-week-old male B6129SF1/J wildtype from Jackson Laboratory. The basic procedure for pupillary light reflex (PLR) recording and pupil size quantification were described in Keenan et al. (*36*). Mice were dark-adapted for 2 hours before the PLR experiment. They were briefly restrained by hand for video recording. Two short videos were shot for each mouse, one for dark-adapted baseline pupil size as a reference point, and the other for pupil constriction measurement after 30 minutes’ light exposure, mainly attributed to melanopsin phototransduction in ipRGCs. A Sony Handycam (FDR-AX33, prime lens, manual focusing, night shot mode, external infrared light source), was used for recording. The dotted pattern from the infrared light source is reflected by the mouse’s cornea and used as the focusing indicator. This setup ensures that the mouse’s pupils can only be focused at a fixed distance from the camera and can be easily visualized in the dark. Light at an intensity of 0.327W/m2 was used to induce PLR. Pupil size was measured as the maximum diameter using Fiji software in each picture frame. The pupil constriction is quantified by normalizing the pupil area at 30 minutes to the baseline area in the dark. A one-way ANOVA statistical analysis was performed.

## Acknowledgement

We are grateful to Drs. Harry Quigley, Elia Duh, Pingwu Zhang and the members of Qian’s lab for their comments and suggestions. We highly appreciate Linda D. Orzolek and Tyler Creamer at the Johns Hopkins Single Cell and Transcriptomics Core for performing the scRNA-seq. We also appreciate the Multiphoton Imaging (MPI) Core in the Department of Neuroscience at Johns Hopkins (5P30NS050274), as well as the Wilmer Eye Institute imaging core (5P30EY001765). We appreciate Yiwen Yan for her participation in quantifying cell survival and the behavioral test. This work was supported by the National Institutes of Health (5R01EY031779 to J. Q. and F. Z.).

## Supplementary Figures

**Fig. S1.**
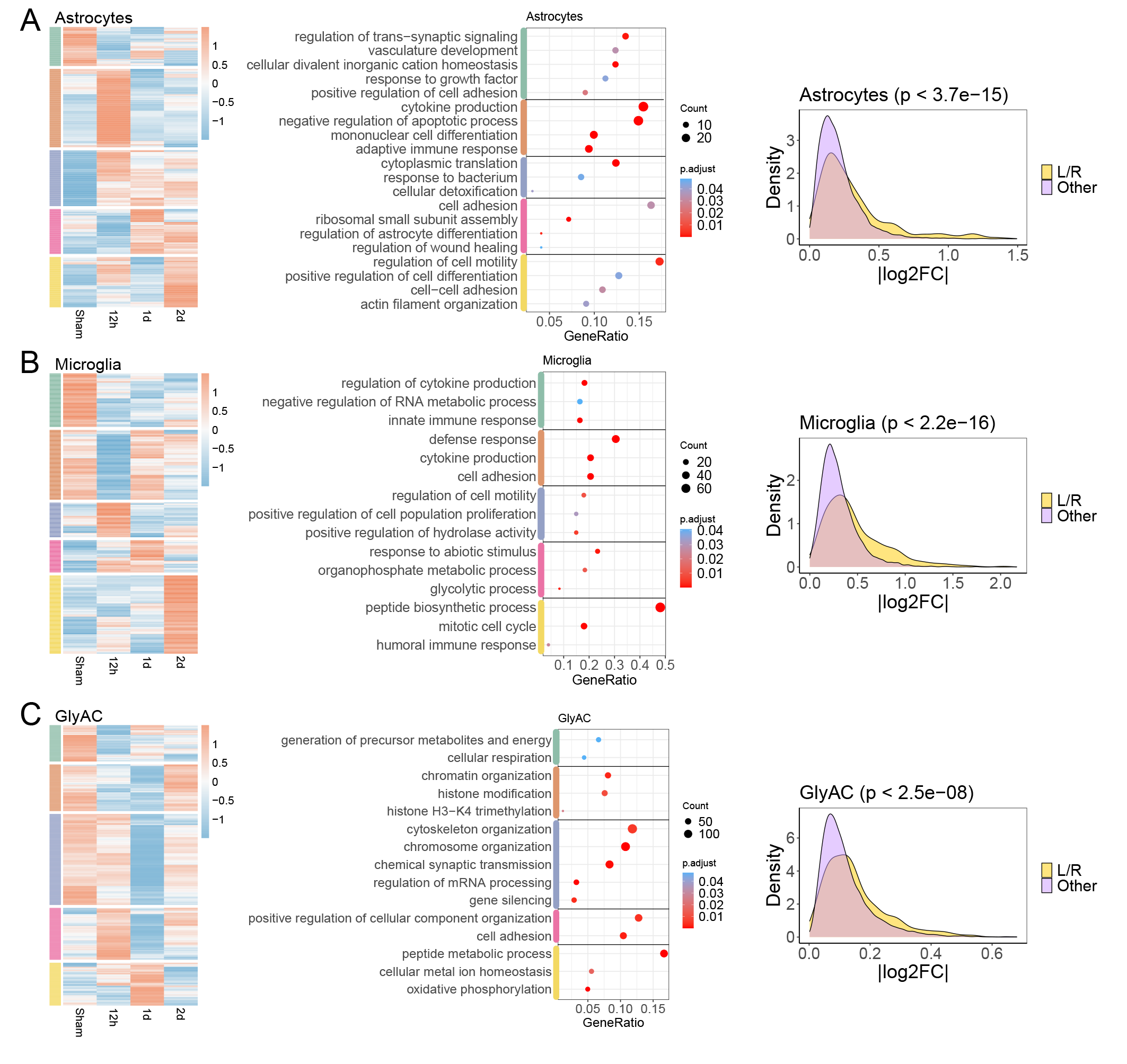
Retinal cells in response to ONC. Response in astrocytes (**A**), microglia (**B**) and glycinergic amacrine cells (**C**), as some examples, after the ONC injury on RGCs. Left panels: DEG heatmaps show dynamical patterns of DEGs at different time points after ONC. Middle panels: Gene ontology analysis reveals representative biological processes enriched by the DEGs in each pattern shown in the heatmaps. Right panel: Density plots of the absolute fold-change (log2FC) values of genes expressed (detection rate > 0.1). Ligand and receptor genes (L/R) shown with yellow color are grouped together, while the other genes are shown with light purple for comparison.

**Fig. S2.**
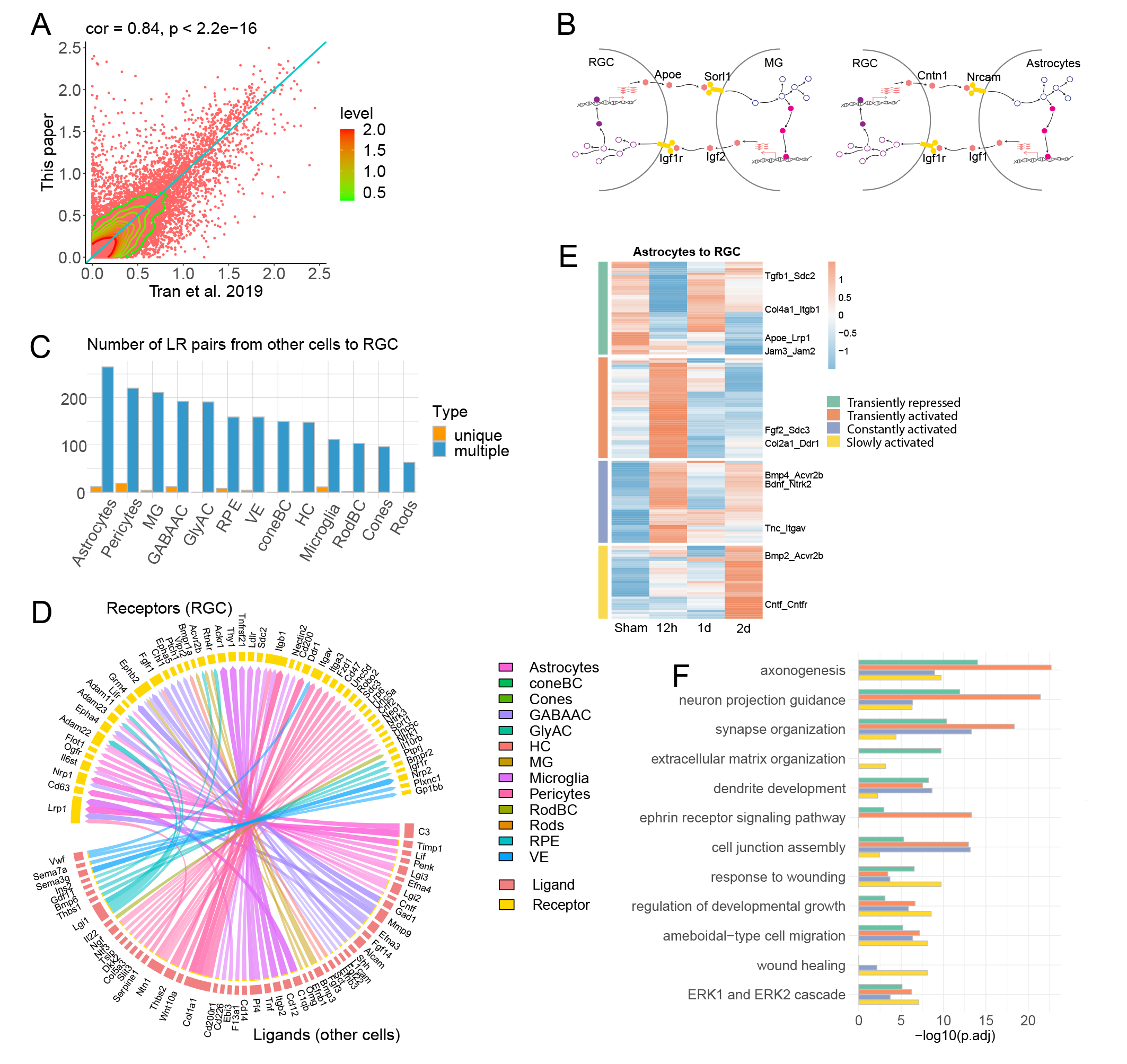
Responsive interactions from other cells to RGCs. (**A**) Correlation of gene expression between RGCs captured and sequenced in whole retina data in this paper and the RGCs in Tran. et al. (**B**) Additional examples of cell-cell feedback loops. The ligand from the sender cell interacts with the receptor on the receiver cell, triggering gene transcription in receiver cells, in which some ligand genes are transcribed and sent back to the original sender cells. (**C**) The number of ligand-receptor interactions from other retinal cells to RGCs. Interactions identified in multiple and unique cell types are colored dark blue and orange, respectively. (**D**) The specific ligand-receptor interactions identified in unique retinal cell type to RGCs, which are shown in orange color columns in the panel (C). Genes in the bottom half of the circle are the ligands secreted from other retinal cells, and the genes in the top half of the circle are the receptors in RGCs. The colors of the connecting edges represent the receiver cell types. (**E**) Heatmap reveals the ONC-induced temporal patterns of interactions from astrocytes to RGCs across time points (the fold change (FC) of the interaction scores between any two time points > 1.2). Four dynamical patterns were identified. (**F**) Gene Ontology analysis (GO) reveals representative biological processes of variable ligand-receptor interactions in patterns identified in panel (E). The enrichment was calculated with all expressed genes in either astrocyte or RGC in any time point as the background (detection rate > 0.1).

**Fig. S3.**
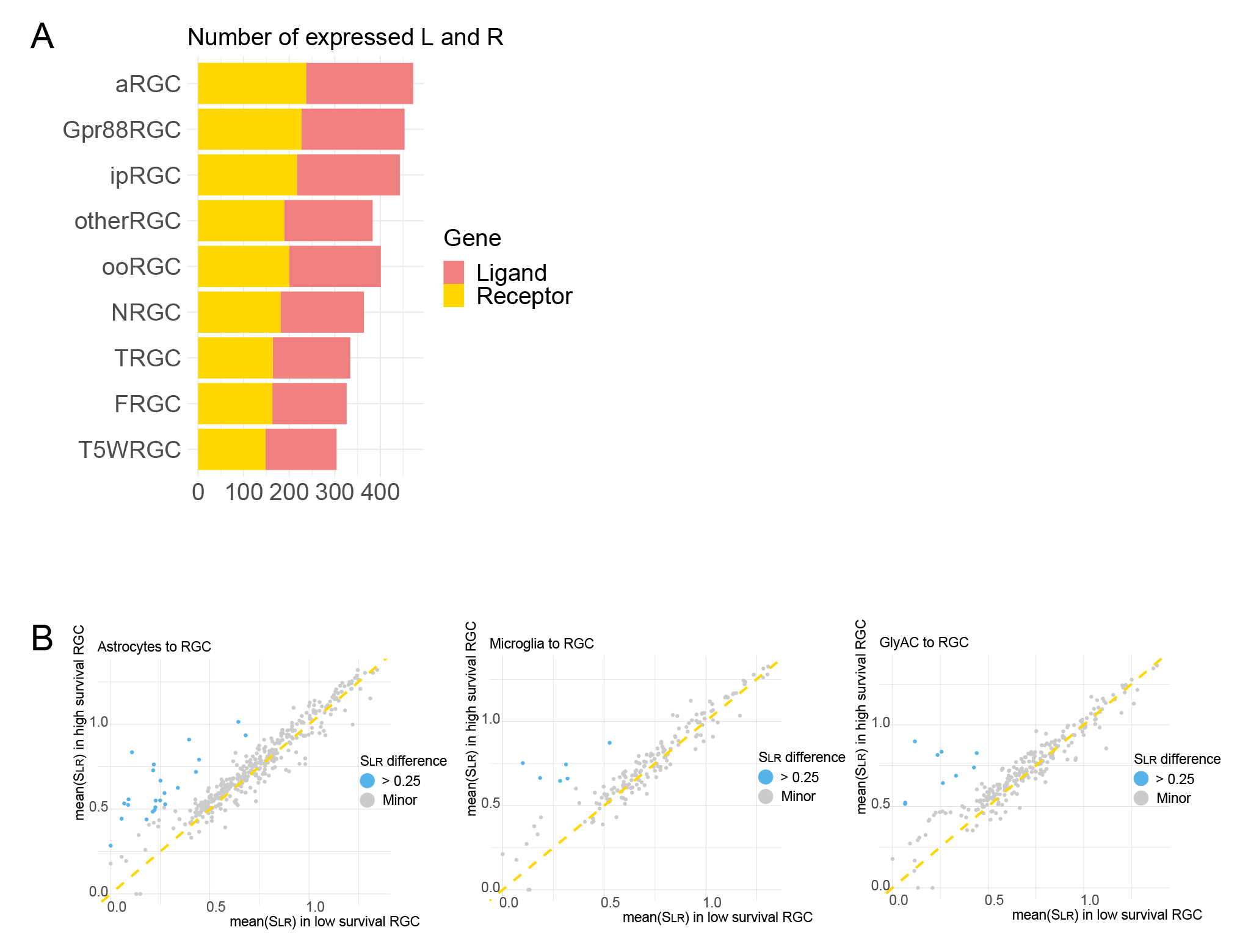
Properties of high-survival vs low-survival RGC subclasses. (**A**) Number of expressed ligand and receptor genes in RGC subclasses (detection rate > 0.1 in any time point). (**B**) Additional examples for calculating protective interactions in astrocytes (left graph), microglia (middle graph) and glycinergic amacrine cells (right graph). For each ligand34 receptor pair, the two calculated mean values of interaction scores, one (y-axis) for the high-and the other (x-axis) for the low-survival RGC subclasses were plotted for comparison. Blue dots are the interactions that are stronger in high-than in low-survival RGC subclasses. Gray dots are the interactions with similar interaction scores between the two categories.

**Fig. S4.**
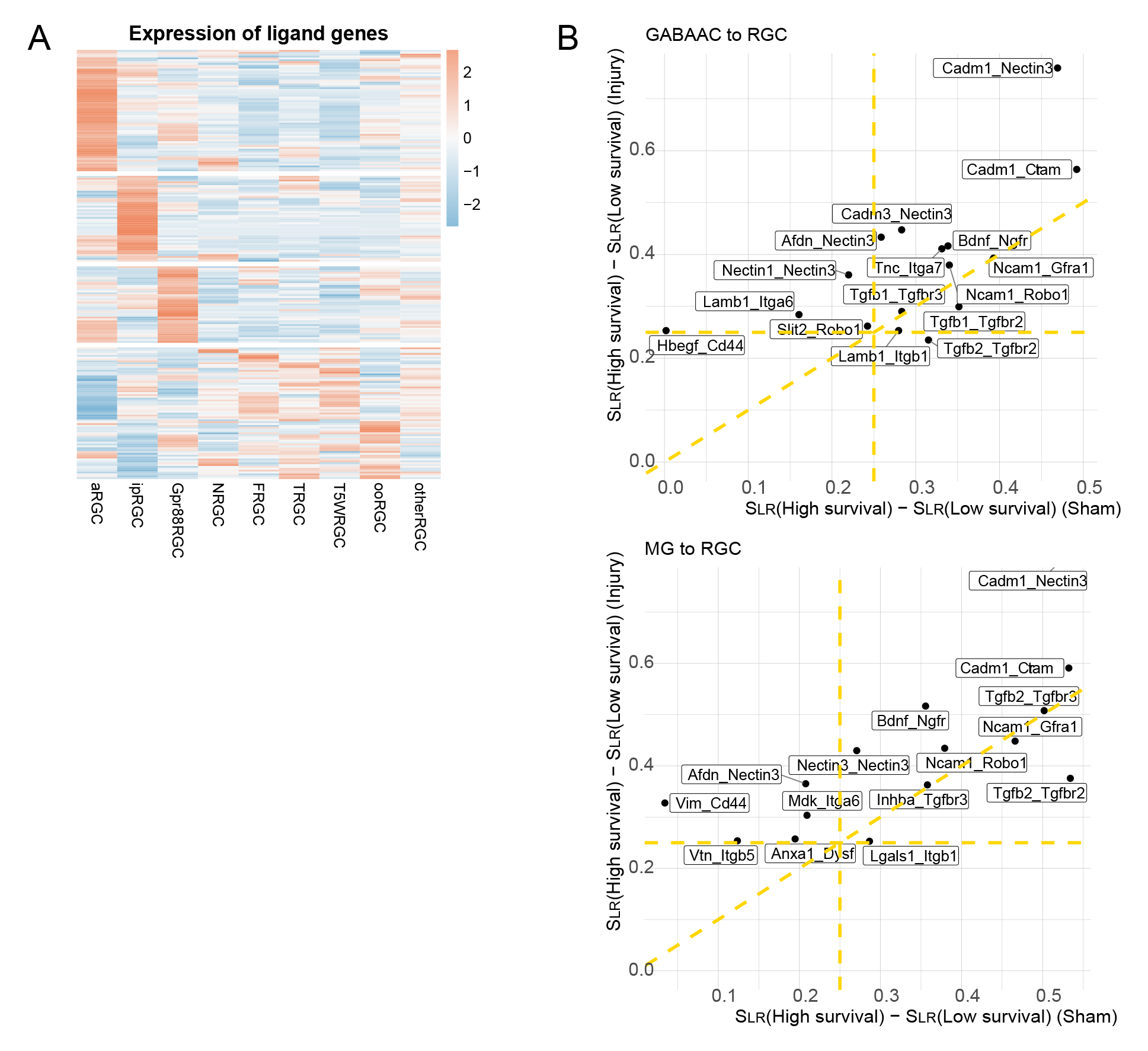
Features of protective interactions. (**A**) Average expression level of ligand genes in RGC subclasses. (**B**) Summary of the preset and induced protective interactions. X-axis is the difference between interaction scores in high-and low-survival subclasses before injury, and Y-axis is the interaction score difference after injury. Interactions from GABAergic amacrine cells to RGCs are shown in upper graph, and the Müller glia to RGCs interactions are shown in lower graph.

**Fig. S5.**
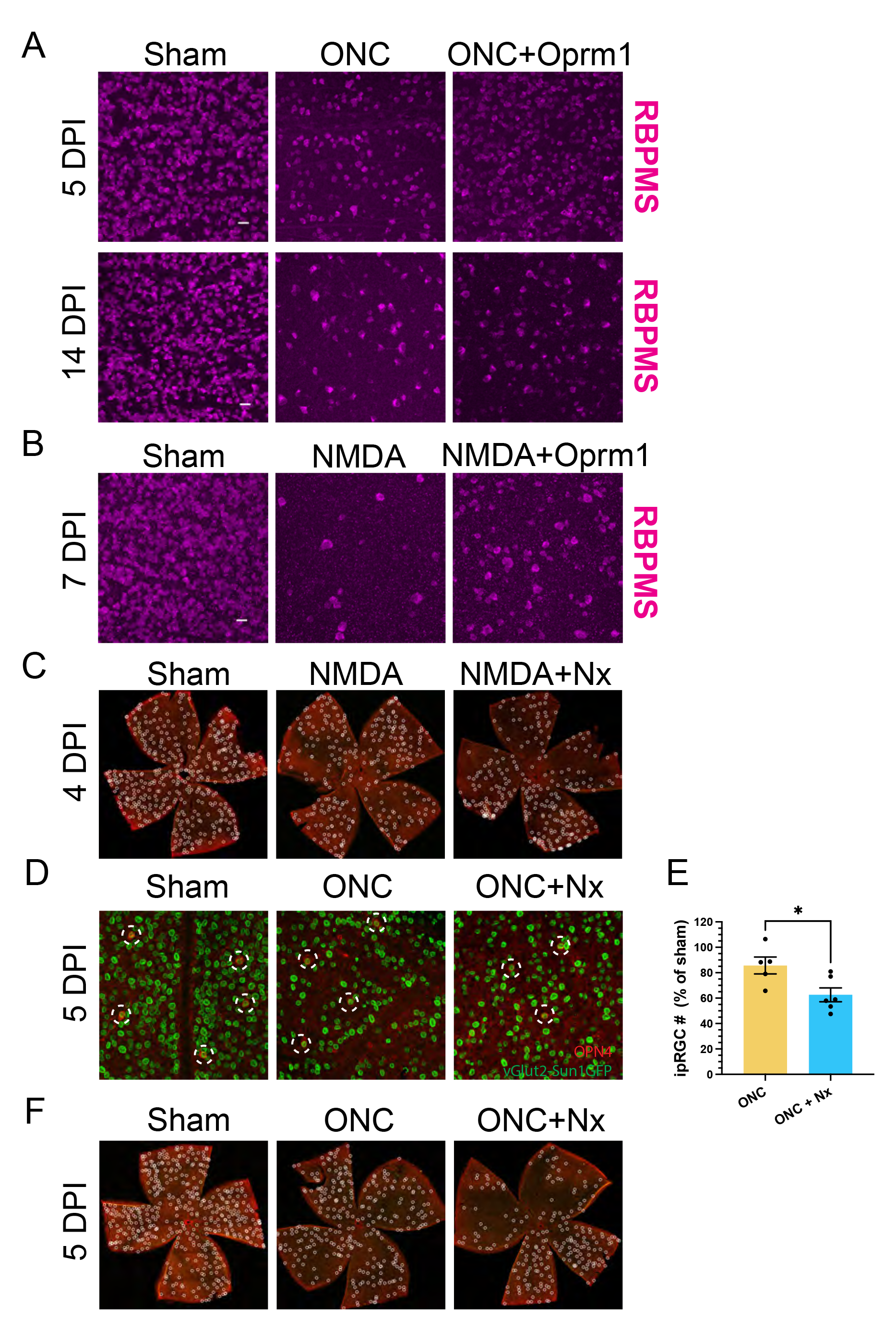
Neuroprotective effect of Oprm1 on RGC survival. (**A**) Representative confocal images of retina whole mounts of RBPMS staining 5 days (upper row) or 14 days (lower row) post-ONC. (**B**) Representative confocal images of retina whole mounts of RBPMS staining 7-days following NMDA damage. (**C**) Representative confocal scans of petal-shape retina whole mounts showing the OPN4 staining of ipRGC, four days after NMDA damage, compared with co-treatment with NMDA and naloxone injection. White dash52 line circles label the ipRGCs (OPN4 in red fluorescence). (**D**) Representative confocal images of retina whole mounts showing ipRGC numbers five days post-ONC. The red channel represents OPN4 staining, and the green fluorescence shows vGlut2-Sun1GFP as pan-RGC marker. White dashed line circles mark the ipRGCs. (**E**) Numbers of ipRGC cell survived (OPN4+), as the percentages relative to the sham group. Data are presented as mean ± SEM. ONC group, n=5; ONC+ Nx group, n=6. One-Way ANOVA, multiple comparison, *p < 0.05. (**F**) Representative confocal scans of petal-shape retina whole mounts showing ipRGC cell survived 5-days post-ONC, and ONC combined with naloxone injection condition. White dashed line circles label the ipRGCs (OPN4 in red fluorescence).

**Fig. S6.**
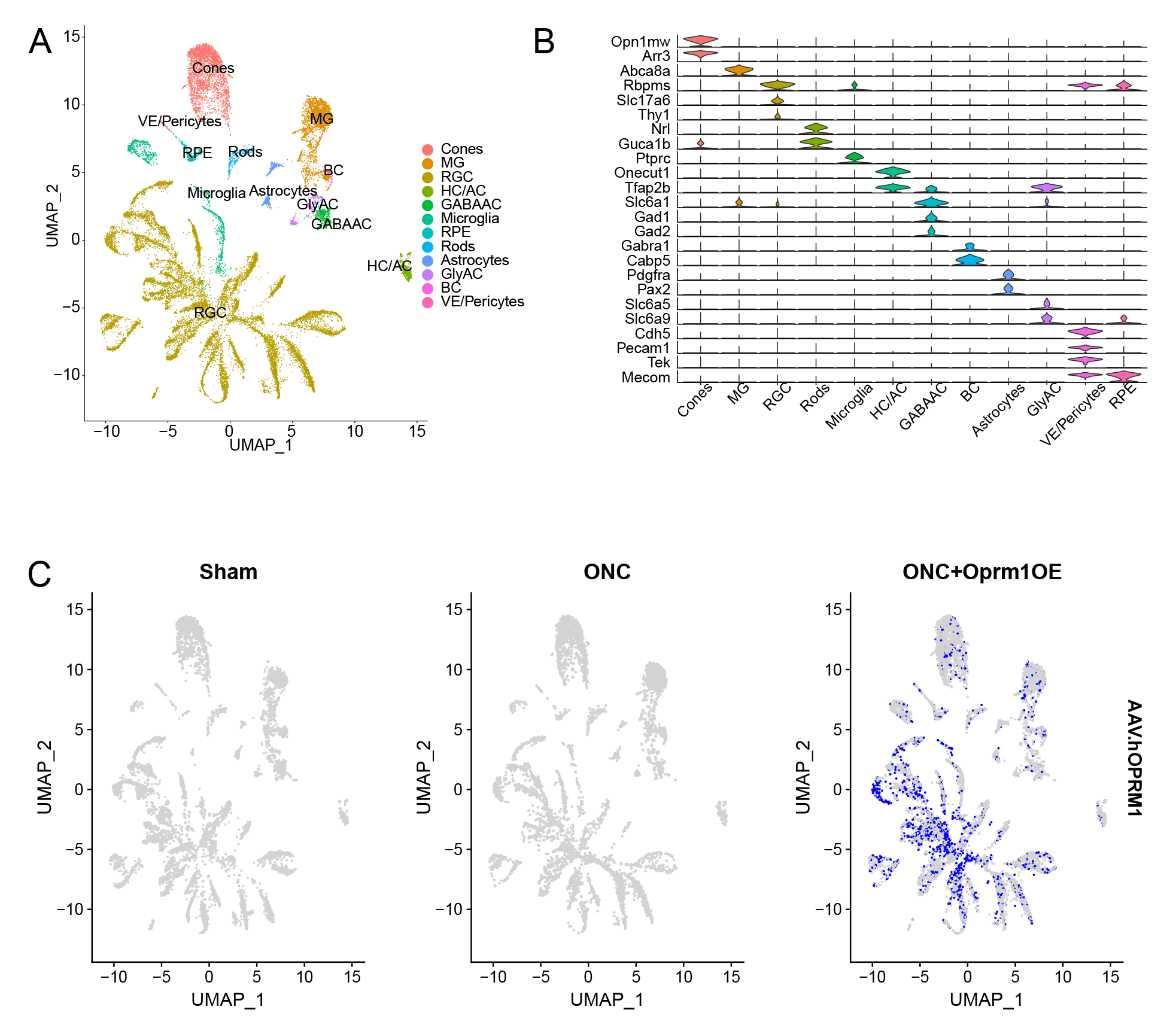
snRNA-seq analysis on sorted RGC nuclei. (**A**) Retinal cells obtained from the samples. The samples were enriched for RGCs with anti-GFP MACS for vGlut2-Sun1GFP+ pan-RGCs. (**B**) Expression level of representative known marker genes in retinal cell types. (**C**) Expression pattens of ectopic human Oprm1 (based on AAV2 WPRE detection) in retinal cells. The ectopic Oprm1 is mainly detected in RGCs.

**Fig. S7.**
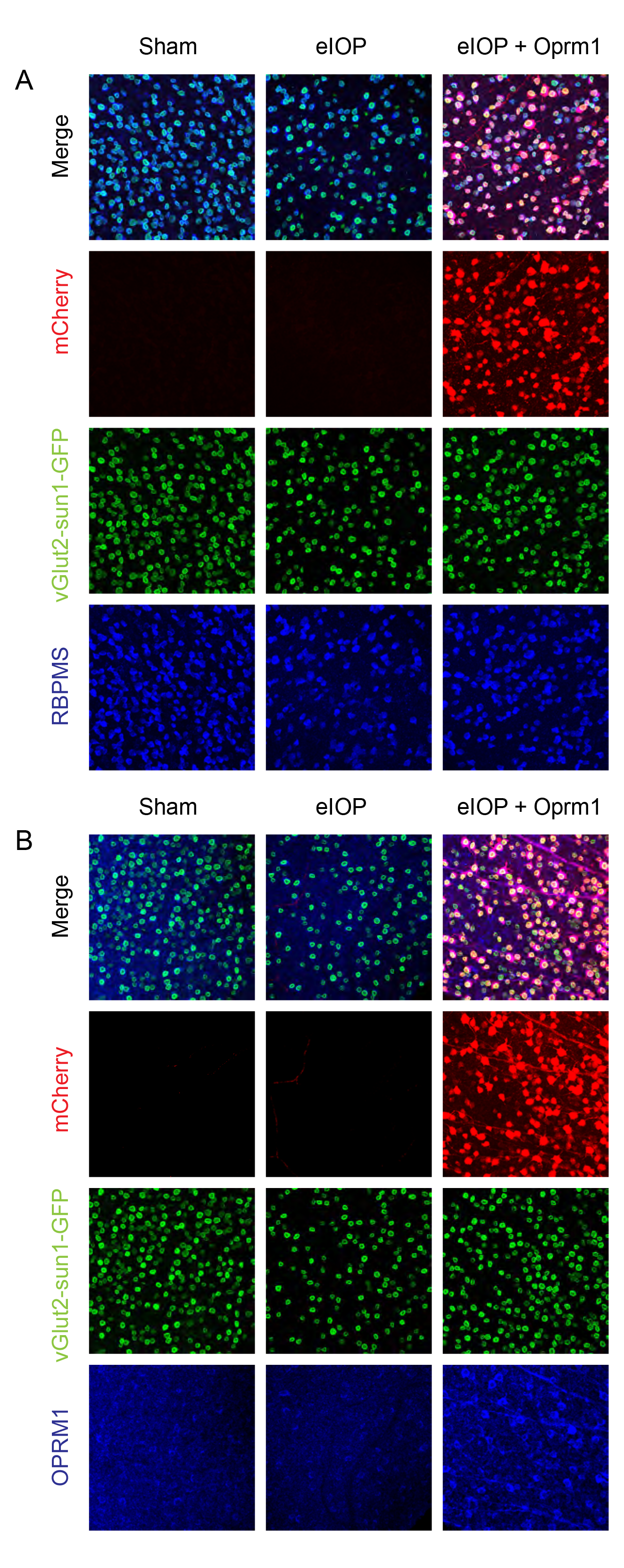
AAV-mediated *Oprm1* gene transduction in eIOP glaucoma model. (**A**) Representative confocal images of retina whole mounts showing the expression of mCherry in Oprm1 + eIOP group, which was transduced with AAV2-FLEX-mCherry-Oprm1 on vGlut2-Cre;LSL-Sun1GFP mice. The red fluorescent staining is mCherry, green fluorescence is Sun1GFP in pan-RGCs, and the blue channel represents RBPMS staining. (**B**) Representative confocal fluorescent images of retina whole mounts showing the expression of mCherry in the Oprm1 + eIOP group, which was transduced with AAV2-FLEX-mCherry-Oprm1, on vGlut2-Cre;LSL-Sun1GFP mice. The red fluorescent staining is mCherry, green fluorescence is Sun1GFP in pan-RGCs, and the blue channel represents Oprm1 staining.

## Notes

### Competing Interest Statement

The authors have declared no competing interest.

